# BAR-CAT: Targeted Recovery of Synthetic Genes via Barcode-Directed CRISPR-dCas9 Enrichment

**DOI:** 10.1101/2025.06.27.658158

**Authors:** Natanya K. Villegas, Mindy H. Tran, Abigail Keller, Calin Plesa

## Abstract

Modern gene-synthesis platforms let us probe protein function and genome biology at unprecedented scale. Yet in large, diverse gene libraries the proportion of error-free constructs decreases with length due to the propagation of oligo synthesis errors. To rescue these rare, error-free molecules we developed BAR-CAT (Barcode-Assisted Retrieval CRISPR-Activated Targeting), an *in-vitro* enrichment method that couples unique PAM-adjacent 20-nt barcodes to each library member and uses multiplexed dCas9-sgRNA complexes to fish out the barcodes corresponding to perfect assemblies. After a single 15-min reaction and optimized wash regime (BAR-CAT v1.0), three low-abundance targets in a 300, 000-member test library were enriched 600-fold, greatly reducing downstream requirements. When applied to 384x and 1, 536x member DropSynth gene libraries, BAR-CAT retrieved up to 122-fold enrichment for 12 targets and revealed practical limits imposed by sgRNA competition and library complexity, which now guide ongoing protocol scaling. By eliminating laborious clone-by-clone validation and working directly on plasmid libraries, BAR-CAT provides a versatile platform for recovering perfect synthetic genes, subsetting large libraries, and ultimately lowering the cost of functional genomics at scale.

## Introduction

Plasmid DNA libraries have been foundational to molecular biology, supporting early gene discovery, physical genome mapping, expression studies, and functional analyses, and serving as a key format in early shotgun genome sequencing (1, 2). Between 2001 and 2021, the cost of sequencing individual human genomes decreased by seven orders of magnitude, outpacing Moore’s Law (3, 4). By 2024, sequencing a human genome costs approximately $200 with the Illumina NovaSeq X, compared to one million USD in 2007 (Illumina). These advances in next-generation sequencing (NGS) technologies shifted the role of plasmid DNA libraries within molecular biology. As a result, plasmid DNA libraries have become essential for large-scale applications, such as barcode lineage tracing (5), CRISPR screens (6), gene regulation studies (7), functional assays of variant effects (8, 9), and directed evolution of enzymes (10).

Modern large-scale applications increasingly rely on synthetic rather than natural sequences. Synthetic libraries are fully customizable and reflect the vast potential design space. Depending on the application, libraries may contain short inserts (<300 bp) or long inserts (>300 bp), with methods tailored accordingly. Short-insert libraries are often generated by microarray or column-based oligo synthesis to encode sequences like peptide fragments, gene exons, DNA barcodes, or guide RNAs. These pools can be PCR-amplified, cloned, or used directly without further assembly. Such libraries have driven progress in functional genomics, for example by enabling pooled CRISPR screens to map enhancer-gene relationships (6), large-scale peptide mapping to identify protein interactions in cancer genes (7), or RNA “bait” libraries for targeted exon sequencing (11).

While synthetic short-insert plasmid DNA libraries have enabled important discoveries, some applications require longer sequences. These include large-scale protein engineering, functional assays of variant effects (MAVEs) (12), synthetic genome assembly, and synthetic metagenomics. Directed evolution experiments often require DNA libraries exceeding 300 nucleotides (nt) to encode full-length protein variants, presenting a challenge for DNA synthesis approaches. Since this length exceeds the limit of standard synthetic oligos, researchers often rely on error-prone PCR (10) or saturation mutagenesis (13) to introduce diversity. Although straightforward, these methods produce variants close to the parental templates, limiting their diversity (13, 14). Synthetic gene libraries offer more control and broader diversity, making them particularly valuable for broad mutation scanning (BMS), a type of MAVE that assesses the functional impact of thousands of homologs across the tree of life. BMS has been used to characterize large libraries of phosphopantetheine adenylyltransferase (PPAT) homologs involved in vitamin B5 biosynthesis (15) and dihydrofolate reductase (DHFR) homologs, uncovering gain-of-function variants that confer resistance to trimethoprim with implications for antimicrobial resistance (16).

DropSynth, a high-throughput gene synthesis method, assembles up to 1, 536 genes in parallel and has made BMS studies feasible by enabling access to large, diverse plasmid DNA libraries (15–17). It assembles user-defined, microarray-derived oligos into full-length genes within emulsion droplets (17), allowing for synthetic gene libraries that are more customizable than those based on natural DNA (15). DropSynth’s scalability is driven by barcoded beads that capture specific oligos. After hybridization, these beads are emulsified with reagents for polymerase cycling assembly (PCA), a PCR-like method that joins overlapping oligos and amplifies full-length genes (18). Emulsion droplets help prevent cross-hybridization between oligos from different gene assemblies. DropSynth reduces gene synthesis costs to approximately US$0.70 per kilobase pair (kbp), compared to the US$70 per kbp cost of traditional synthesis methods.

Currently, DropSynth gene assemblies are limited to around 1, 000 base pairs (bp), with only about 8% of constructs perfectly assembled at the DNA level (19). This limitation stems from errors introduced during phosphoramidite oligo synthesis (POS), originally developed by Beaucage and Caruthers, which accumulates truncations, deletions, and substitutions as oligos grow longer (4, 20, 21). POS uses a four-step cycle with roughly 99% efficiency per nt addition (21), but as oligo length increases, yields decline and error rates rise (22). As a result, assembling longer genes, such as those 2, 000 bp in length, likely yields less than 1% perfect constructs due to compounded oligo synthesis errors and additional errors during polymerase cycling assembly in emulsions (19). MAVEs and other multiplex functional assays perform better with higher rates of perfect assemblies. Identifying perfect constructs with very low representation (<1%) demands substantial oversampling, increasing costs (4). While commercial vendors can synthesize oligos up to 300 nt with per-nucleotide error rates as low as 1 in 2, 000 to 1 in 3, 000 nt (vendor-provided rates), the amount of perfect full-length products decreases sharply with length due to the cumulative effect of errors and reduced synthesis yield.

To address these challenges, several strategies have aimed to enrich error-free oligos during or after synthesis. One approach uses sequencing-by-synthesis (SBS) to extend oligos hybridized to universal primers until they reach the correct length, at which point a biotinylated reversible terminator is incorporated to isolate full-length products (23). Other methods leverage next-generation sequencing (NGS) to select validated oligos before assembly. For example, an earlier approach used Roche 454 sequencing with robotic retrieval of error-free oligos (24), but this has been replaced by more accurate and affordable Illumina platforms (25). To improve scalability, Schwartz and colleagues developed dial-out PCR, which adds unique adaptors to oligo pools, sequences them, and selectively amplifies perfect sequences (26). Though effective, this method is expensive and difficult to scale (4, 22). Hybridization-based enrichment offers better scalability by melting away mismatched oligos during annealing. However, this method only modestly improves error rates, such as to 1 in 1, 394 bp (27), and requires carefully designed probes to avoid capturing erroneous sequences (28). It also lacks the resolution to discriminate highly similar sequences (11).

To enable functional studies of longer genes using DropSynth and other assembly methods, it is crucial to enrich perfect full-length gene assemblies directly. A traditional approach involves cloning and validating assembled genes by Sanger sequencing (4, 29). While accurate, this method validates only one gene at a time and is labor-intensive and costly (30, 31). Nanopore sequencing enables validation of gene sequences across multiple plasmids simultaneously (32), but each plasmid must still be individually isolated for retrieval, limiting scalability. Dial-out PCR can also retrieve perfect genes or oligos (26, 33), but requires independent reactions and custom primers for each targeted gene, limiting scalability.

More scalable strategies for enriching perfect gene assemblies include fusing fluorescent or antibiotic resistance markers, enabling functional selection. Assemblies with premature stop codons fail to express the marker and are removed (34), though re-initiation at downstream start codons can yield truncated proteins, leading to false positives (35). Another approach uses MutS, a mismatch repair protein that binds mismatched base pairs in heteroduplex DNA (36). Beads or columns functionalized with MutS can enrich perfect assemblies up to 25.2-fold (4, 37), though this requires large amounts of heteroduplex input DNA (17, 38). Other mismatch-recognizing enzymes like T7 endonuclease I, used for perfect gene selection also require large amounts of intact heteroduplexes and do not cleave all mismatch types, as shown by persistent errors in Sanger sequencing (39). The low quantity of heteroduplex DNA produced by multiplex gene assembly methods like DropSynth limits the application of error-correcting enzymes such as MutS and T7 endonuclease I.

Given the lack of sufficiently targeted, affordable, and scalable methods for enriching perfect gene assemblies from synthetic gene libraries, we sought to develop a solution that could not only address this gap but also have broader applications. This method must be sufficiently versatile to enrich DNA sequences beyond perfect gene assemblies, such as selectively enriching specific gene subsets from large gene libraries. This flexibility would help save time, labor, and resources. In addition to its immediate utility for DropSynth, we envisioned it as a tool with potential applications in emerging fields such as DNA data storage (40), offering a “cut and paste” mechanism for large-scale DNA manipulation.

We selected *Streptococcus pyogenes* Cas9 (SpyCas9), a well-characterized nuclease from the CRISPR gene-editing suite, as the core of our DNA enrichment method. CRISPR-Cas9 is ideal for this application due to its precision, scalability, and ability to multiplex with minimal cross-talk among targets, all driven by programmable single-guide RNAs (sgRNAs) (41–43). Its extensive characterization and broad compatibility with various tools make it a reliable choice for diverse DNA manipulation applications (44).

To minimize off-target effects associated with CRISPR-Cas9, we chose the catalytically inactive version, dCas9, which offers reduced leakiness. In a CRISPR-based high school education kit, dCas9:sgRNA ribonucleoprotein (RNP) complexes repressed the expression of chromoproteins eforRed and fwYellow 88-fold and 10-fold more, respectively, than active Cas9 RNPs (45). While the magnitude of repression varied between chromoproteins, dCas9 consistently reduced the leakiness of gene repression, which is a key feature for improving target specificity. The versatility of dCas9 is further demonstrated by its successful use in a cell-free diagnostic test to enrich rare alleles, illustrating its potential for multiplex DNA enrichment *in vitro* (43). These features make dCas9 an ideal candidate for precise, multiplexed DNA manipulation in our approach.

Here, we introduce Barcode-Assisted Retrieval – CRISPR-Activated Targeting version 1.0 (BAR-CAT v1.0), an *in vitro* method for selectively retrieving barcoded genes and their corresponding plasmids from DNA plasmid libraries. BAR-CAT v1.0 requires DNA libraries to be tagged with random 20-nt PAM-adjacent barcodes using a barcoding plasmid. These barcodes serve as binding sites for dCas9 complexed with sgRNAs containing matching spacers. After sequencing of the barcoded libraries, barcodes are mapped to perfect gene assemblies or other DNA sequences of interest (**Fig. 1A**). A custom computational pipeline filters spacers that may lead to dCas9 off-target binding and organizes the remaining spacers into sub-libraries. These sub-libraries are transcribed into sgRNA libraries each targeting specific gene libraries, and synthesized *in vitro* with T7 RNA polymerase (T7 RNAP) in a single-pot reaction (46). The sgRNAs are incubated with biotinylated dCas9 to form RNPs, which are then pulled down using streptavidin-coated magnetic beads. Non-target sequences are washed away, and the enriched genes are PCR-amplified, sequenced, and analyzed to compare barcode frequencies before and after enrichment (**Fig. 1A**).

**Fig. 1.**
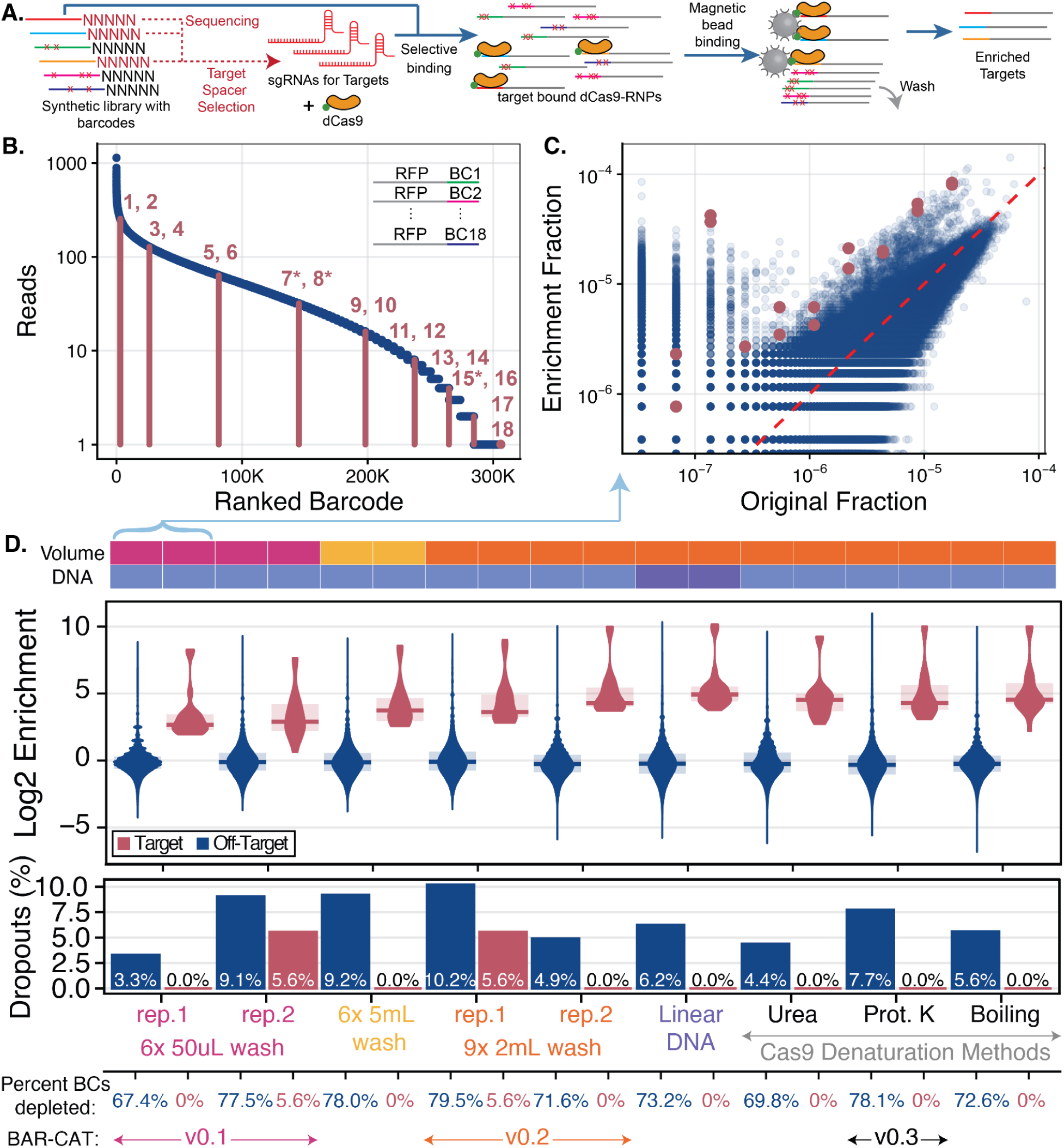
Overall workflow for BAR-CAT proof-of-concept and initial optimizations. **A.** A synthetic gene library is first generated and sequenced to map 20-nt barcodes to their corresponding DNA molecules. Barcodes associated with perfect sequences are selected as protospacers for targeting. Spacer sequences are then *in vitro* transcribed into sgRNA libraries, which are complexed with dCas9 to form ribonucleoprotein complexes (RNPs). These RNPs are incubated with the synthetic library, allowing dCas9 to bind targeted barcodes. Biotinylated dCas9 is captured using streptavidin-coated magnetic beads, followed by bead pulldown and washes to remove unbound, non-target sequences. Enriched DNA is PCR-amplified and sequenced via nanopore to assess target enrichment. **B.** Rank-ordered read abundance of 18 selected target barcodes from a >300, 000-barcode single-gene *rfp* library. Each dot represents a barcode, ranked by read count. Dark magenta lines denote target barcodes, with magenta numbers indicating barcode IDs. Asterisks mark the three target barcodes used in BAR-CAT optimizations shown in **Fig. 2**. Scatter plot comparing the fraction of barcode reads in the *rfp* library before (original) and after enrichment. Each blue dot represents a non-target barcode; each magenta dot represents a target barcode. The red dashed unity line denotes equal representation before and after enrichment, serving as a reference for assessing enrichment magnitude. **D**. Enrichment conditions are shown chronologically along the x-axis. The colored bar above the plots summarizes key iterative protocol changes, including bead wash volume, DNA input format (linear vs. supercoiled), and dCas9 denaturation method. Violin plots display the distribution of log_2_ enrichment scores for off-target (blue) and target (magenta) barcodes. Shaded areas represent the interquartile range (25th–75th percentile), with horizontal bars indicating median enrichment. Percent dropouts are indicated in bar plots for off-target (blue) and target (magenta) barcodes. All barcodes with reduced abundance (log_2_ enrichment < 0), including barcodes that drop out post-enrichment, are reported as % depleted. The pink condition (6× 50 μL) corresponds to the original proof-of-concept protocol (v0.1) and is linked to panel **C** by a blue arrow.

To make BAR-CAT v1.0 scalable and multiplexable, we aimed to identify and extend the limits of CRISPR-dCas9 multiplexed binding. Multiplexed CRISPR editing enables simultaneous regulation of multiple loci, facilitating studies of complex phenotypes and genetic interactions (47–49). In mammalian cells, CRISPR-Cas9 has been used for multiplex knockouts of up to three targets per cell, but scaling beyond this is limited by the need to express each sgRNA from its own promoter, complicating cloning and delivery (48, 50). In contrast, cell-free systems like BAR-CAT v1.0 bypass this limitation by using *in vitro* transcribed sgRNAs added directly to CRISPR reactions, enabling more extensive multiplexing. Cell-free CRISPR-dCas9 with *in vitro* transcribed sgRNAs has shown success in diagnostic applications for enriching rare alleles (43, 51). Thus, the development of BAR-CAT v1.0 provided an opportunity to examine the constraints and parameters of CRISPR-dCas9 binding *in vitro*, informing future optimizations and expanding the potential of multiplexed CRISPR systems.

In this study, we demonstrate that optimizations to BAR-CAT, including increased bead washes and dCas9 denaturation prior to amplification, modestly improve enrichment of 18 targeted barcodes by approximately 3-fold. The most robust improvement (median ∼600-fold for three barcodes) was achieved by increasing input gene library amounts tenfold, likely improving the DNA to dCas9 ratio and reducing off-target enrichment. When applied to DropSynth-assembled dihydrofolate reductase (DHFR) libraries containing 384 or 1, 536 genes, we observed up to 149-fold enrichment for 12 targets. However, at larger library sizes, enrichment became inconsistent, and target dropout rates increased. These results suggest further optimization strategies and provide insights for scaling BAR-CAT and other *in vitro* CRISPR methods to improve multiplexing and reduce off-target effects.

## Results and Discussion

### Proof-of-concept enrichment of 18 barcodes from a barcoded mCherry library (BAR-CAT v0.1)

We sought to test whether the BAR-CAT method could enrich specific barcodes of interest from a low-complexity DNA library prior to scaling up to more complex DropSynth gene libraries. To this end, we constructed a barcoded red fluorescent protein (*rfp)* gene library by PCR-amplifying the *mCherry* gene using in-house primers and cloning it directly adjacent to 20 nucleotide (nt) unique barcodes within a barcoding plasmid, pEVBC3. This plasmid was derived from the pEVBC vector previously used to barcode DropSynth gene assemblies (15), but lacked a PAM site adjacent to the 3′ end of the barcode on the template strand, which is required for CRISPR-Cas9 to initiate R-loop formation on the non-template strand. This omission was intentional, as our goal was to rapidly develop a proof-of-concept using a single-gene library. The barcoded *rfp* library contained sufficient barcode diversity to select targets that naturally contained a 5′-NGG-3′ PAM sequence at the 3’ end of the variable barcode region. These targets were used for initial proof-of-concept experiments and subsequent optimization.

Following a standard workflow, we ligated *mCherry* amplicons into pEVBC3 to produce a 2.7 kb barcoded *rfp* plasmid, transformed it at high efficiency into *E. coli* DH5α, pooled the resulting colonies, isolated plasmid DNA, and sequenced the library on an Illumina MiSeq to quantify barcode diversity for BAR-CAT target selection (**Fig. 1A**). Sequencing revealed >300, 000 unique barcodes, comparable in diversity to DropSynth gene libraries. Barcode representation varied widely, ranging from approximately 1, 000 reads for the most abundant barcodes to 1 read for the least. This variability can be summarized using the Gini Coefficient, a metric of inequality where a value of zero corresponds to perfect uniformity and a value of one to perfect inequality (19, 52). The observed Gini Coefficient of 0.56 highlights the skewed distribution typical of synthetic DNA libraries, where biases introduced during oligonucleotide synthesis, amplification, assembly, or cloning can result in significant disparities in sequence abundance (53).

In contrast, natural genomes have relatively uniform sequence composition. For example, the least-represented target in our synthetic *rfp* library was detected as a single read among 14, 641, 735 total reads, corresponding to 6.8 × 10^-8^ of all molecules. By comparison, a single-copy gene in diploid human genomic DNA occurs in two copies per 6.2 Gb genome, which corresponds to approximately 3.2 × 10^-6^ of all molecules (assuming a 10 kb gene) and 47-fold more abundant than our scarcest target. Whereas human genomic DNA is essentially uniform in copy number, our synthetic library exhibited far greater inequality (Gini: 0.56). This highlights the need for targeted enrichment methods like BAR-CAT to compensate for representation biases within synthetic gene libraries, particularly when downstream applications depend on uniform sequence recovery.

To minimize these abundance-dependent biases during enrichment, we selected 18 target barcodes spanning this range. Two barcodes were chosen per read abundance tier, from the highest (barcodes 1 and 2, ∼1, 000 reads) to the lowest (barcodes 17 and 18, 1 read), producing a balanced set for this initial proof-of-concept (**Fig. 1B**). These barcodes were used as protospacers for CRISPR-dCas9 targeting (**Fig. 1A**).

To generate the corresponding sgRNA library, synthetic DNA oligos encoding the 20 nt spacer sequences were ordered with flanking PCR priming sites and a BsaI type IIS restriction site. Oligos were pooled and assembled via Golden Gate assembly (GGA) to produce DNA templates for *in vitro* transcription (IVT). The resulting pooled template was transcribed by T7 RNAP into an 18-plex sgRNA library, which was validated by gel electrophoresis and RNA-seq (**Fig. 1A**). Our full IVT protocol and validation results are detailed elsewhere (46).

For targeted dCas9 enrichment, we selected the dCas9-3XFLAG™-Biotin protein (Sigma). After RNPs form with this biotinylated dCas9 protein and the18-plex sgRNA library, dCas9 should selectively bind to the 18 targeted barcodes. We reasoned that streptavidin magnetic beads would be a simple, effective, and straightforward way to capture the biotinylated dCas9 (**Fig. 1A**), as demonstrated by other CRISPR-Cas9 targeted DNA enrichment methods (51).

After the RNPs have bound to barcode targets following enrichment, streptavidin-coated magnetic beads were incubated with the biotinylated RNP-DNA complexes. Subsequent washes were performed using a wash buffer to remove non-specifically bound DNA, thereby isolating the biotinylated complexes retained on the beads (**Fig. 1A**). To identify optimal washing conditions for enriching targeted barcodes with high sensitivity, we screened four buffer compositions for wash stringency. A sham DNA capture experiment was performed using a synthetic library containing a single barcode from the *rfp* library. This DNA was incubated with equilibrated streptavidin beads in the absence of biotinylated dCas9 or RNPs, simulating nonspecific DNA-bead interactions (**Fig. S1A**). Beads were washed either three or six times with 50 µL of each buffer, and wash steps were carried out at either room temperature (RT, ∼25 °C) or 30 °C to assess temperature effects on wash stringency.

After the final wash, supernatants were subjected to quantitative PCR alongside a 10 ng positive control and a no-template control. Amplification cycle (Cq) values were plotted to evaluate wash efficiency, where higher Cq values indicated reduced residual DNA and thus greater stringency (**Fig. S1A**).

The buffer conditions tested were: immobilization buffer (IB), 2× binding and wash buffer (2× B&W), 2× B&W supplemented with 10% NP-40, and 1× TE supplemented with 10% NP-40 (**Table S1**). The 2× B&W buffer, containing 2 M NaCl in TE with EDTA, is recommended by the bead manufacturer. The NP-40-supplemented variant was included to test whether the non-ionic detergent would enhance stringency. In contrast, IB lacked salt but included DTT to stabilize dCas9, while the 1× TE + NP-40 formulation combined detergent without salt.

We did not observe any differences among wash buffer stringency between the two temperatures tested (RT or 30 °C) or between conditions using three versus six 50 µL washes.When data were combined across temperatures and wash numbers, the 2× B&W (21.5 ± 1.12, n = 6) and 2× B&W + 10% NP-40 (22.0 ± 0.48, n = 4) wash conditions exhibited mean Cq values that were 20.1% and 18.2% lower than the no-template control (NTC; 26.9 ± 4.07, n = 6), respectively. In comparison, IB (17.4 ± 0.69, n = 3) and 1× TE + 10% NP-40 (15.4, n = 1) conditions showed even lower values (more DNA), with mean Cq values 35.5% and 42.9% lower than the NTC, respectively. Altogether, these results suggested that 2× B&W and 2× B&W + 10% NP-40 were the most stringent washes tested (**Fig. S1B**). Due to limited sample size and overlapping trends, we could not definitively distinguish between the two, so 2× B&W was selected for all subsequent BAR-CAT experiments. This buffer offered both reliable performance and ease of preparation, and its selection aligned with the manufacturer’s recommendations.

We first aimed to use our 18-plex sgRNA library to enrich the 18 corresponding barcode protospacers from the *rfp* library. To form active RNPs, we combined 3 µL of 33 nM sgRNA library with 1 µL of 1 µM biotinylated dCas9, maintaining a 1.11:1 dCas9-to-sgRNA molar ratio, in a total volume of 27 µL, following NEB recommendations. Complexes were pre-incubated for 10 minutes at 25 °C, followed by an additional 10-minute incubation at 37 °C. Then, 50 ng of the *rfp* plasmid library was added as a binding substrate, corresponding to a 39.6:1 RNP-to-DNA molar ratio, and incubated for 15 minutes at 37 °C to allow selective binding.

We added sufficient streptavidin magnetic beads (5 µL), capable of capturing 25 pmol of biotinylated dCas9, providing approximately 25-fold excess relative to the dCas9 input. Enrichment reactions were incubated with the beads for 30 minutes at 37 °C with shaking. The beads were then washed six times with 50 µL of 2× B&W buffer, a condition previously determined to minimize non-specific retention of off-target barcodes. qPCR was performed to determine the optimal number of cycles to avoid overamplification, followed by PCR amplification of enriched barcodes directly from the beads for nanopore sequencing.

Analysis of the sequencing data showed that all 18 targeted barcodes were more abundant after enrichment compared to the original *rfp plasmid library* (magenta dots), indicating successful enrichment. This is evident from their position above the red dashed unity line, which marks equal abundance in both libraries (**Fig. 1C**). However, two distinct subpopulations of off-target barcodes (blue dots) were observed: (1) a majority that remained at moderate abundance (10^-5^ to 10^-6^ fraction), and (2) a smaller group that were unexpectedly enriched, with representation similar to that of the targeted barcodes. The first subpopulation was unexpected, as we expected fewer off-target barcodes after enrichment. This is because washing the streptavidin-coated magnetic beads should have removed those not specifically bound to biotinylated dCas9:target barcode complexes. In contrast, we attributed the second subpopulation, consisting of enriched low-abundance off-target barcodes (10^-8^ to 10^-6^ fraction), to known off-target binding of CRISPR-dCas9 (54–56). We identified that errors arising during sgRNA spacer oligo synthesis, PCR amplification (2.8 × 10^-7^ errors/nt), and IVT by T7 RNAP (5 × 10^-5^ errors/nt) (57) resulted in production of some mutant sgRNA spacers within the 18-plex library (46). These mutations could have contributed to off-target binding if present at the PAM distal region at the 5’ end of the sgRNA spacers, known to tolerate mismatches to targets (56).

Overall, the enrichment of 18 targeted barcodes was moderately successful, establishing an initial proof of concept for the BAR-CAT design. Log_2_ enrichment scores, defined as the log_2_ fold change in barcode abundance before and after enrichment, are shown in **Fig. 1D** for replicate 1 of the 6 × 50 µL bead wash condition (shown in pink). This protocol was designated as BAR-CAT version 0.1 (v0.1). Most off-target barcodes had log_2_ enrichment scores near zero, indicating no enrichment. Barcodes with scores between −6 and 0 were considered depleted relative to the input library. Barcodes with no sequencing reads (scores below −6) were classified as post-enrichment dropouts (**Fig. 1D**).

We observed that 67.4% of off-target barcodes were depleted, and 3.3% completely dropped out following enrichment, suggesting that the BAR-CAT v0.1 bead washing conditions were insufficient to fully eliminate off-target sequences. In contrast, the 18 targeted barcodes were enriched with a median log_2_ fold change of 2.7 (6.3-fold enrichment), with no evidence of target depletion. To assess the practical impact of this enrichment, we calculated the population fraction enrichment, defined as the ratio of the sum of the population fractions of all target barcodes after enrichment to the sum of their fractions before enrichment. This metric yielded a 6.4-fold increase, indicating that approximately 6.5-fold fewer colonies would need to be screened to recover clones bearing the desired barcodes.

### Bead wash optimization and development of BAR-CAT v0.2

Although the initial post-enrichment results were promising, they exhibited a high level of non-enriched, off-target barcodes, indicating insufficient removal of unwanted sequences during the bead wash steps. High-stringency washes are critical in hybridization-based DNA capture methods for eliminating cross-hybridized and unbound sequences (11). Drawing from this precedent, we hypothesized that increasing the number and volume of bead washes would help reduce the retention of non-target barcodes caused by weak or non-specific dCas9 interactions in the BAR-CAT protocol. Having previously optimized wash buffer stringency (**Fig. S1)**, we next tested whether increasing wash volume and frequency could further enhance the removal of off-target barcodes.

We conducted a preliminary experiment to assess the appropriate number of washes. To do this, we incubated 50 ng of supercoiled *rfp* library DNA with magnetic SNAP-Capture bead aliquots, using these beads as a proxy for evaluating wash efficiency (not as a replacement for the streptavidin magnetic beads). The beads were washed 15 times with 1 mL of IB. Supernatants from washes 3, 6, 9, 12, and 15, along with the beads from the final wash, were collected for quantitative PCR (qPCR) and gel electrophoresis to quantify the abundance of the 586 bp *rfp* library PCR product.

For this preliminary experiment, we used IB rather than the 2× B&W buffer normally used in BAR-CAT washes, due to the known inhibitory effects of high salt on PCR (58, 59). The PCR product band became undetectable in the wash supernatant after the ninth wash, while PCR of beads washed 15 times still produced abundant product (**Fig. S2A**). Quantification of the PCR product gel bands using ImageJ (60) revealed a DNA peak intensity of 18, 552 a.u. upon amplification of residual DNA from the supernatant of the third wash. In comparison, 69.23% and 78.41% less DNA was detected in washes 9 (5, 708 a.u.) and 12 (4, 005 a.u.), respectively (**Fig. S2B**). Based on these results, we increased the cumulative 2X B&W buffer washes of streptavidin beads to 30 mL (6x 5 mL) and 18 mL (9x 2 mL). The 18-plex sgRNA library and protocol described for BAR-CAT v0.1 were used, with the only variation being the increased number of washes. Additionally, we included an enrichment control representing BAR-CAT v0.1 washed with a cumulative wash volume of 0.3 mL (6x 50 µL, replicate 2).

For replicate 2 of the original 6× 50 µL bead wash condition, the barcode distribution closely matched that of replicate 1 (**Fig. 1C)**, confirming the reproducibility of our experimental control (**Fig. S3A**). In contrast, the 6× 5 mL (**Fig. S3C**) and 9× 2 mL (**Fig. S3B**) bead wash conditions displayed a noticeable shift in target barcode enrichment compared to the original distribution. A key difference was observed in the distribution of off-target barcodes: a greater number of moderately abundant off-target barcodes from the original population (fractional abundance between 10^-5^ and 10^-6^; **Fig. 1C**, **Fig. S3A**) were enriched in these higher-volume wash conditions, likely due to a reduction in background signal from non-enriched barcodes (**Fig. S3B, C**). We hypothesized that this apparent enrichment reflected increased dCas9 off-target binding that had been obscured in the 6× 50 µL control due to higher background levels. These findings suggest that increasing bead wash volume can reduce background DNA and enable finer resolution of enrichment dynamics.

The control 6× 50 µL condition (replicate 2, shown in pink) yielded a median log_2_ enrichment of 2.9 for the 18 targeted barcodes, corresponding to a 7.4-fold increase, with a population enrichment score of 5.8. These values closely matched those observed in replicate 1, confirming protocol reproducibility. However, 1 of the targeted barcodes was depleted relative to the input library, and one barcode dropped out (5.6% target dropouts) entirely post-enrichment, indicating reduced recovery for certain targets. Among off-target barcodes, 78% were depleted and 9.9% dropped out, representing a 14% increase in depletion and a threefold increase in dropouts compared to replicate 1 (**Fig. 1D**). These results suggest that while overall enrichment performance was reproducible, some stochastic variation in barcode retention may still occur between replicates.

The 6× 5 mL bead wash condition (shown in yellow) produced a median log_2_ enrichment of 3.7 of 18 targeted barcodes (12.5-fold enrichment) and a 13.4-fold population fraction enrichment with no depletion or dropout of targets. The 9× 2 mL condition (orange, replicate 1) showed a median log_2_ enrichment of 3.6 (12.2-fold increase) and a 12-fold population fraction enrichment, but with 5.6% target depletion and 5.6% target dropouts. Similar percentages of off-target barcodes dropped out with increased washes compared to the 6× 50 µL wash control (replicate 2). Specifically, the 6× 5 mL wash condition yielded 78.0% depleted off-target barcodes and 9.2% dropouts, while the 9× 2 mL wash condition (replicate 1) showed 79.5% depletion and 10.2% dropouts. (**Fig. 1D**). Overall, increasing the cumulative wash volume by 100-fold (6× 5 mL) and 60-fold (9× 2 mL) resulted in 2.3- and 2.1-fold improvements in target barcode enrichment, respectively. Based on these results, we selected the 9× 2 mL bead wash condition for all subsequent enrichments and designated this protocol as BAR-CAT v0.2, as it provided better enrichment with less buffer usage and labor compared to the 6× 5 mL condition.

### Approaches to denature dCas9 and reduce off-target barcodes for development of BAR-CAT v0.3

While optimizing bead washes yielded modest improvements, we sought to further increase enrichment of targeted barcodes and reduce off-target carryover. We hypothesized that denaturing dCas9 post-enrichment would release bound targets while reducing off-target barcode retention by removing beads and dCas9 complexes. Additionally, we tested whether using a linearized version of the *rfp* library could enhance enrichment by potentially enabling dCas9 to more efficiently access target PAM sites.

We tested these conditions experimentally as follows. Eighteen targets were enriched from either the supercoiled or linearized *rfp* library using the 18-plex sgRNA pool. The beads used to capture dCas9 for these conditions were washed according to the BAR-CAT v0.2 protocol (9× 2 mL), and three dCas9 denaturation methods were tested for their ability to release bound DNA: 8 M urea (43), proteinase K (61, 62), or heat incubation at 65 °C for 5 minutes (62, 63). Following urea or proteinase K treatment, supernatants containing liberated DNA were collected, column-cleaned, PCR-amplified, and nanopore-sequenced to identify enriched barcodes. The linear *rfp* library template (827 bp) was generated by PCR amplifying the supercoiled plasmid using primers *mi9_FWD_amp_NV* and *mi9_REV_amp_NV* (**Table S2**) for 14 cycles. An important distinction between the linear and supercoiled *rfp* plasmid libraries is the effect of molecule length on the RNP to DNA ratio. Although the same masses of DNA, sgRNA, and dCas9 were used for both enrichments, maintaining an RNP to target ratio of 1.11, the shorter length of the linear *rfp* library (827 bp) resulted in approximately 3.5 times more DNA molecules. This increase in DNA copy number reduced the overall RNP to DNA molar ratio from 35 to 1 in the supercoiled condition to 10 to 1 in the linear condition.

The distributions of target and off-target barcodes across the tested conditions were similar to those observed in the 9× 2 mL replicate 1 condition (orange) (**Fig. S3B**; see also **Fig. S4A–E**). This indicates that, at a global level, neither the use of a linear *rfp* library nor dCas9 denaturation appreciably altered the representation of target or off-target barcodes. This outcome contrasts with the substantial differences observed between the 6× 50 µL (**Fig. 1C**, **Fig. S3A**) and 9× 2 mL (**Fig. S3B**) bead wash conditions, where wash stringency had a pronounced effect on barcode distributions.

Next, we examined the log_2_ enrichment scores for the linear and supercoiled *rfp* plasmid library conditions. The supercoiled *rfp* library control (9 × 2 mL, replicate 2, shown in orange) yielded a median log_2_ 4.3-fold enrichment of the targeted barcodes (19.5-fold enrichment) and 23-fold population fraction enrichment. Replicate 2 of BAR-CAT v0.2 (9 × 2 mL) showed a 1.6-fold higher median enrichment than replicate 1 of the same condition, possibly due to experimental variation. Enrichment of targets from the linear *rfp* library yielded a median log_2_ 4.9-fold enrichment (31-fold enrichment) and 26-fold population fraction enrichment. This represented a 1.6-fold increase in median target enrichment relative to the supercoiled condition (9 × 2 mL, replicate 2). While this difference could be attributed to the linear DNA format or experimental variation, it is also possible that the increased molar concentration of DNA molecules in the linear library due to their shorter length led to a slight increase in targeted enrichment.

We then examined the log_2_ enrichment scores for dCas9 denaturation using urea, proteinase K, and heat. All three methods produced similar targeted enrichment values compared to those observed with the supercoiled and linear *rfp* libraries. Urea treatment produced a median log_2_ 4.5-fold enrichment of targeted barcodes (23-fold enrichment), with 14-fold population fraction enrichment. Proteinase K treatment resulted in a median log_2_ enrichment of 4.3 (20.0-fold enrichment), with 22-fold population fraction enrichment. Lastly, boiling yielded a median log_2_ enrichment of 4.5 (23-fold enrichment), with 22-fold population fraction enrichment. None of the tested dCas9 denaturation enrichments showed depletion or dropout of targeted barcodes (**Fig. 1D**).

To evaluate the impact of the tested conditions on off-target barcode representation, we compared depletion and dropout rates across conditions. The supercoiled plasmid DNA enrichment control showed 71.6% depletion and 4% dropouts (83, 414 barcodes). The linear DNA enrichment yielded a slight increase in both metrics, with 73.2% depletion and 6% dropouts (106, 119 barcodes), potentially reflecting experimental variation or enhanced depletion due to the linear format. Urea-mediated dCas9 denaturation resulted in lower depletion (69.8%) and 4.4% dropouts (74, 542 barcodes) compared to the supercoiled and linear conditions. Boiling produced slightly higher depletion (72.6%) and 5.6% dropouts (95, 186 barcodes). The highest depletion (78.1%) and dropout rate (7.7%, 131, 254 barcodes) were observed with proteinase K treatment, representing a 3.7% increase in depletion compared to the supercoiled control (**Fig. S5**, **Fig. 1D**). Despite this, proteinase K did not alter enrichment scores for targeted barcodes relative to the supercoiled and linear conditions, suggesting it effectively removed off-targets while preserving on-target signals. We concluded that urea treatment was the least effective, likely due to its harsh, non-specific denaturation mechanism. In contrast, proteinase K offers an enzymatic means of dCas9 denaturation, which may limit the release of non-specifically bound off-target barcodes into the supernatant. By comparison, urea and boiling treatments may promote the dissociation of both enriched and non-specifically bound DNA, reducing overall enrichment quality.

Based on these findings, we adopted proteinase K-mediated dCas9 denaturation for all subsequent enrichments and designated this protocol optimization as BAR-CAT v0.3. Although the linearized *rfp* library contained roughly 3.5-fold more DNA molecules than the supercoiled control and accordingly showed a modestly higher enrichment of the 18 targets, the reason for this improvement remains unclear. It may result from increased DNA copy number, differences in DNA conformation, or experimental variation.

### Assessing DNA input amount and incubation time on enrichment performance to develop BAR-CAT v1.0

Since targeting 18 barcodes from the linearized *rfp* library resulted in a modest increase in enrichment compared to the supercoiled version (**Fig. 1D**), we considered several possible explanations. One possibility is that Cas9 binds more efficiently and specifically to relaxed, linearized DNA than to the negatively supercoiled DNA such as that of our synthetic *rfp* plasmid library. This idea is supported by a previous study showing that Cas9 exhibits reduced off-target binding and cleavage on relaxed DNA substrates compared to negatively supercoiled DNA (64). However, our linear DNA enrichment condition also included 3.5 times more DNA molecules than the supercoiled condition, introducing a confounding variable that may have contributed to the observed increase in median log_2_ enrichment scores (**Fig. 1D**). Therefore, we hypothesize that the improved enrichment observed with linear DNA was primarily driven by the greater molar abundance of DNA molecules available for dCas9 binding.

Supporting this hypothesis is the observation that targeted constructs in our library are far less abundant than genomic regions enriched by previously published methods, such as RNA-guided endonuclease (RGEN) enrichment methods such as RGEN-R and RGEN-D (62). Although genomic DNA is more uniformly represented than synthetic DNA libraries, RGEN enrichment methods require 10–20 μg of fragmented DNA per reaction, suggesting that high input DNA is important for effective enrichment. In RGEN-R, Cas9 cleaves large genomic fragments (>20 kbp), which are then captured via biotinylated adaptors and streptavidin beads. In contrast, RGEN-D employs dCas9 pre-complexed with biotinylated sgRNAs for direct pulldown, followed by proteinase K treatment to release the DNA. These workflows rely on over 200-fold more DNA than our standard BAR-CAT reactions (62). Together, these comparisons support the idea that improved enrichment with linear *rfp* DNA is at least partly due to increased molar abundance of targetable molecules, which is especially important given the low representation and unequal distribution of targets in synthetic libraries.

Based on these observations, we hypothesized that increasing the amount of supercoiled *rfp* plasmid library DNA would enhance the fold enrichment of targeted barcodes. To test this, we enriched three barcodes (7, 8, and 15) from the supercoiled *rfp* library (barcodes with asterisks, **Fig. 1B**) using two DNA input amounts: the original 50 ng and a 10-fold higher input of 500 ng. The 500 ng input DNA condition decreased the ratio of RNPs to DNA to approximately 4:1 from the 40:1 ratio used previously (50 ng input DNA). We targeted three barcodes for this experiment due to their consistent enrichment performance across conditions when 18 barcodes were targeted. More specifically, barcodes 7 and 8 showed modest enrichment in the BAR-CAT v0.3 enrichment condition (proteinase K denaturation, **Fig. 1D**) while barcode 15 was one of the two most enriched targets of all 18 barcodes (**Fig. S6**). To streamline the experiment, we used synthetic sgRNAs (Integrated DNA Technologies) due to concurrent challenges with optimizing our IVT sgRNA workflow (46).

We retained targeted enrichment of the *rfp* library with supercoiled plasmid topology for several reasons. Although the linearized version showed slightly better enrichment, it had been prepared using 14 cycles of PCR, raising concerns about potential bias. PCR can skew barcode representation by preferentially amplifying GC-rich sequences and introducing mutations in spacer regions, which may increase off-target dCas9 binding (65–67). Since BAR-CAT already includes a post-enrichment PCR step, we sought to minimize additional amplification. Restriction digestion could, in principle, offer a PCR-free linearization strategy. However, it would complicate the workflow and, importantly, many enzymes would cleave within the 20 bp random barcodes, removing or truncating a portion of uniquely identifiable molecules.

We also faced practical constraints on DNA concentration, as input DNA needed to be concentrated to ≤3 µL to maintain consistent reaction conditions. Using column-based purification, we were typically able to concentrate the supercoiled *rfp* plasmid library to ∼167 ng/µL, making 500 ng inputs feasible. However, higher concentrations were difficult to achieve reliably with our existing protocol, so we capped the input at 500 ng. While actual concentrations varied between preparations, this amount represented the practical upper limit. Future BAR-CAT optimizations could incorporate tRNA-assisted ethanol precipitation to further concentrate DNA, potentially enabling >5 µg of input DNA in ≤3 µL. This approach has been shown to enhance the concentration of genomic DNA libraries and improve plasmid transformation efficiency by up to 400-fold compared to conventional methods (68).

In addition to testing 500 ng and 50 ng DNA input amounts, we evaluated three enrichment incubation times at 37 °C: 15 minutes (our original condition), 1 hour, and 8 hours. The 15-minute time was based on NEB recommendations, but other Cas9 enrichment methods have various incubation times. RGEN-D uses 20 minutes (62), Aalipour and colleagues used 30 minutes (43), Kim and colleagues used 2 hours (51), and RGEN-R used 8 hours (62). While the rationale behind these incubation times is not always specified, *in vitro* Cas9 binding studies provide some context. A Cas9 binding rate constant of 0.8 ± 0.2 min^-1^ has been reported (63), implying that ∼80% of target DNA can be bound within one minute under simplified conditions with a single target on a plasmid. However, BAR-CAT employs a highly multiplexed library with ∼300, 000 barcodes, which increases search complexity. We therefore hypothesized that longer incubation times might improve enrichment by giving dCas9 more time to locate targets. To test this, we included 1- and 8-hour incubation times to assess whether extended search duration would enhance recovery of the three selected barcodes.

All three targeted barcodes showed substantial increases in log_2_ enrichment scores across all tested DNA input amounts and incubation times, with no target dropouts observed. (**Fig. 1D**). The 50 ng input DNA and 15 min incubation control condition (BAR-CAT v0.3) produced a median log_2_ 6.1-fold enrichment (70-fold enrichment) and 135-fold population enrichment of three targets (**Fig. 2A**). This represents a 3.6-fold increase in median enrichment compared to the corresponding 18-plex condition with proteinase K treatment (**Fig. 1D**), suggesting that dCas9 binding is more efficient with fewer targets, resulting in stronger signals. However, it may also reflect the effect of targeting three well-performing barcodes, which could inflate enrichment scores by skewing the median upward. Nevertheless, the 500 ng input with 15-minute incubation condition yielded a median log_2_ enrichment of 9.2 (600-fold enrichment) and a 1, 094-fold population enrichment. This corresponds to an 8.6-fold increase in median enrichment compared to the 50 ng input (**Fig. 2A**).

**Fig. 2.**
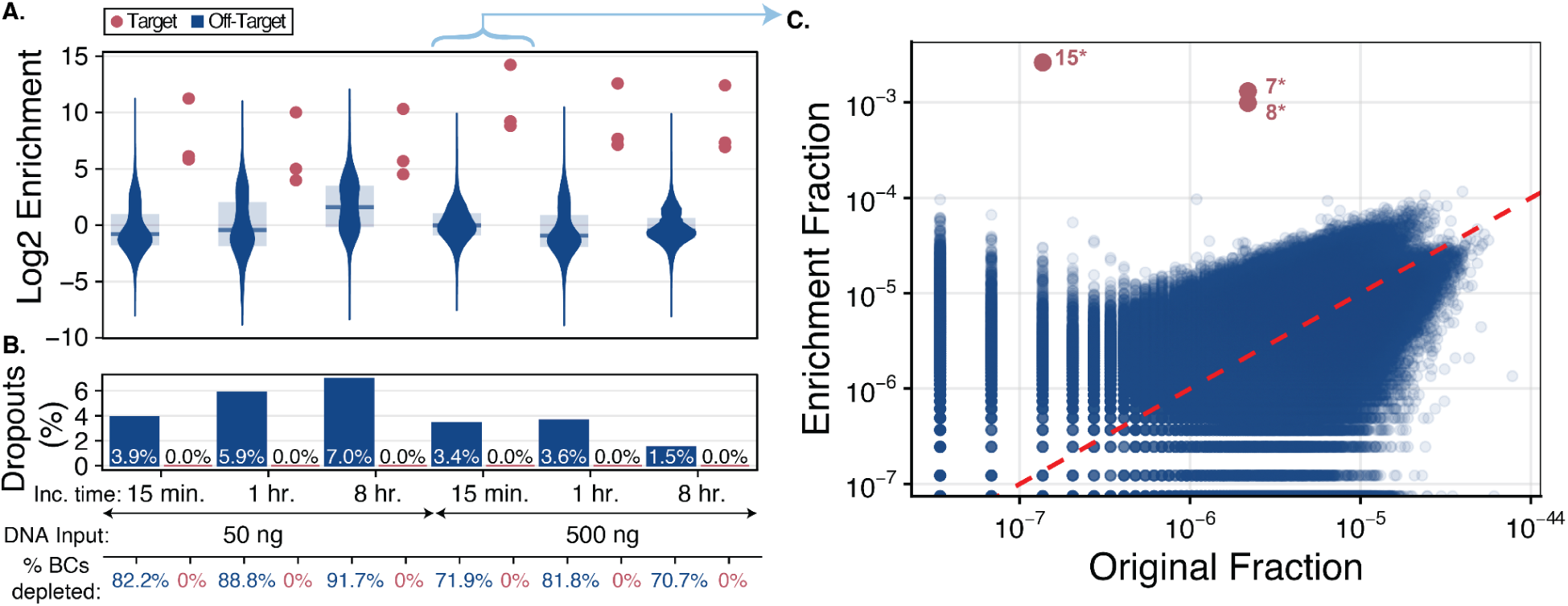
Increasing the input supercoiled *rfp* library DNA leads to large improvements in observed BAR-CAT enrichment while increasing incubation time does not. **A.** Violin plots showing log_2_ enrichment scores, calculated as the log₂ fold change in barcode abundance before and after enrichment. Each distribution compares non-target (blue) and target (magenta) barcodes across conditions varying in enrichment incubation time and supercoiled *rfp* library DNA input amount. Shaded regions represent the interquartile range (25th–75th percentile) for non-target barcodes, with bars indicating median log₂ enrichment. The condition using 500 ng of input library DNA and a 15 min incubation corresponds to the optimized BAR-CAT protocol (v1.0) and is linked to panel **C** by a blue arrow. The best-performing enrichment condition (15 min incubation, 500 ng input DNA) resulted in a median log₂ enrichment of 9.2 (600-fold) and a 1, 094-fold population fraction enrichment. This result indicates that increasing the input library DNA amount from 50 ng to 500 ng improves BAR-CAT enrichment. Conversely, maintaining a short enrichment time (15 min) increases median log_2_ enrichment values while increasing incubation time to 1 or 8 hours leads to slight decreases in enrichment. **B**. Percent dropout values are listed in the bar plots for target (magenta) and non-target (blue) barcodes. Protocol variables tested, such as enrichment incubation time and DNA input amount, are indicated below the dropout bars. Barcodes with reduced abundance (log₂ enrichment < 0), including dropouts, are reported as % BCs depleted. **C**. Scatter plot comparing the fraction of barcode reads in the *rfp* library before (original) and after enrichment for the best performing enrichment condition (15 min, 500 ng input DNA) shown in panel **A**, as indicated by the blue arrow. This condition, referred to as BAR-CAT v1.0, was used for all subsequent experiments. Blue dots represent non-target barcodes; magenta dots represent target barcodes. Asterisks denote the three protospacers selected from the original set of 18 target barcodes (**Fig. 1B**). The red dashed unity line marks equal representation before and after enrichment and serves as a reference for evaluating enrichment.

Next, we evaluated the percentages of off-target barcode depletion and dropout across these conditions. Both metrics were similar between the 50 ng, 15 min incubation (82.2% depletion, 3.9% dropouts) and 500 ng, 15 min incubation (79.1% depletion, 3.4% dropouts) conditions (**Fig. 2A**). The distribution of target and off-target barcodes before and after enrichment with 500 ng DNA for 15 minutes showed that off-target barcodes aligned closely with the dashed red unity line, indicating that nearly all off-targets were non-enriched (**Fig. 2C**, **Fig. S7D**). In contrast, the 50 ng DNA condition displayed a distinct population of moderately enriched off-target barcodes **(Fig. S7A**). These results suggest that increasing the DNA input amount reduces off-target enrichment, improving CRISPR-dCas9 binding specificity. However, it did not eliminate off-target, non-enriched barcodes, which are potentially a result of the magnetic bead pull-down and washing approach overall (**Fig. 2C**).

We then evaluated enrichment scores for the 50 ng and 500 ng supercoiled *rfp* plasmid library DNA input amounts incubated for 1 hour and 8 hours. The 50 ng condition showed decreased target enrichment at these longer incubations compared to the 15-minute incubation. The 1-hour enrichment produced a median log₂ 5.0-fold enrichment (32-fold), a 2.2-fold decrease relative to the 15-minute time point. The population enrichment score similarly decreased from 135-fold to 55-fold (2.5-fold decrease). For the 8-hour enrichment, the median log₂ 5.7-fold (57-fold) enrichment represented a 1.6-fold increase compared to the 1-hour incubation, while the population score increased by 1.4-fold (55 to 75-fold). The 500 ng conditions showed similar trends. The 500 ng, 1-hour enrichment condition had a median log₂ 7.7-fold enrichment (205-fold), roughly 2.9-fold lower than the 500 ng, 15-minute incubation, while the population score decreased 3.1-fold (1, 094 to 354-fold). The 8-hour 500 ng incubation yielded a median log₂ 7.4-fold enrichment, similar to the 1-hour time point, with a slight decrease in population score by 1.3-fold (205 to 163-fold) (**Fig. 2A**). None of the three targets were depleted or dropped out in any condition (**Fig. 2B**). Overall, these results show that targeted enrichment decreases after 1 hour of incubation within the tested DNA input range.

We then evaluated the off-target barcode depletion and dropouts across incubation times for the 50 ng supercoiled *rfp* plasmid library DNA input enrichments. For the 50 ng input conditions, the 1 hour (88.8% depletion, 5.9% dropouts) and 8 hour (91.7% depleted, 7.0% dropouts) increased slightly compared to the 15 min incubation condition (82.8% depleted, 3.9% dropouts (**Fig. 2A**). Surprisingly, while the distribution of target and off-target barcodes appeared similar between the 15 min (**Fig. S7A**) and 1 hour incubation enrichments **(Fig. S7B**), the 8 hour condition showed that the off-target, non-enriched barcode population had decreased (**Fig. S7C)**. Since the 8 hour condition had a slightly higher sequencing depth compared to the others [data not shown], we concluded that the observed decrease in non-enriched barcodes was likely due to the longer incubation time. However, since the log_2_ fold enrichment was lower for the 8 hour condition (**Fig. 2A**) compared to the other incubation times with 50 ng input DNA, we concluded that increased depletion of off-target barcodes did not improve targeted enrichment.

Next, we evaluated whether similar trends in off-target barcode depletion and dropouts observed for the 50 ng DNA inputs were comparable to those observed across incubation times for the 500 ng DNA inputs. We did not observe a substantial increase in off-target depletion for the 1-hour (81.8% depletion, 5.9% dropouts) or 8-hour (70.7% depletion, 1.5% dropouts) incubations compared to the 15-minute incubation (71.9% depletion, 3.4% dropouts) (**Fig. 2A**). Meanwhile, the distribution of target and off-target barcodes was similar for the 15 min (**Fig. S7D**), 1 hour (**Fig. S7E**), and 8 hour incubation enrichments with 500 ng input DNA (**Fig. S7F**). Based on these results, we observed decreased depletion in the 8 hour incubation time with 500 ng of input DNA compared to 15 min and 1 hr incubations with 500 ng input DNA. This is the opposite of the trend observed for incubation time with 50 ng of input DNA.

Overall, these results show that the 500 ng input DNA condition yields superior enrichment of the three targeted barcodes across all incubation times compared to 50 ng. However, 1 hour of incubation reduced median enrichment by 2.2-fold (50 ng DNA) and 2.9-fold (500 ng DNA) relative to the 15-minute incubation. This was unexpected, since approximately 97% of Cas9 is estimated to be bound to target DNA at one hour of incubation based on cleavage assays (63, 69), although these estimates may not fully reflect dCas9 binding dynamics over longer timescales.

One possible explanation is that a fraction of dCas9 molecules dissociate and fail to rebind due to sgRNA degradation, which can occur after 8 hours at 37 °C (70). Another potential explanation is that the chemically synthesized sgRNAs used in these experiments may contain synthesis errors. These sgRNAs were purified by standard desalting, and IDT estimates an error rate of ∼0.05 errors per 100 nt. Errors within the reversibility determining region (RDR; nucleotides 8–17), in particular, are known to promote dCas9 dissociation when mismatches occur in this region (56). Therefore, dCas9 molecules complexed with synthetic sgRNAs harboring errors in the RDR may initially bind their targets but dissociate over time, although we have not experimentally verified this in our system.

While the reason for reduced enrichment at longer incubation times remains unclear, we concluded that a 15-minute incubation with 500 ng input DNA provided the optimal conditions tested. Accordingly, these parameters were incorporated into BAR-CAT v1.0, which was then used for scale-up evaluation of perfect gene assemblies from DropSynth gene libraries.

### Performance of BAR-CAT v1.0 in the Scale-Up of Target Enrichment from DropSynth Libraries

Given the promising performance of BAR-CAT v0.1 in enriching three barcodes from the supercoiled *rfp* plasmid library using synthetic sgRNAs, we applied this method to enrich perfect gene assemblies from two DropSynth dihydrofolate reductase (DHFR) gene libraries. One library contained 1, 536 DHFR genes (495–579 bp; library S2), while the other contained 384 DHFR genes (504–576 bp; library S4). These libraries were previously assembled in-house for a study investigating mutations in DHFR homologs that confer trimethoprim resistance (16).

To enable barcode targeting, DHFR inserts were cloned into a pEVBC3-derived plasmid (pEVBC8), modified to include a 5′-AGG-3′ protospacer adjacent motif (PAM) at the 3’ end of each barcode. This modification was necessary because DropSynth libraries yield relatively few perfect assemblies per target, limiting our ability to computationally select barcodes that end in a GG dinucleotide, as we had done with the *rfp* library and the original pEVBC3 backbone, which lacked a PAM site.

Barcoding was performed by ligating DHFR products into barcoded pEVBC8 using T4 DNA ligase, as done for the *rfp* library. The resulting plasmids ranged from 2.5 to 2.57 kbp. We transformed the plasmid libraries into electrocompetent 10-beta *Escherichia coli*, then isolated plasmid DNA for Illumina MiSeq sequencing to assess barcode coverage and guide sgRNA selection. We recovered 5, 300, 938 unique barcodes from the 384-gene library, and 93, 418 from the 1, 536-gene library.

Perfect DHFR gene assemblies were mapped to barcodes using a custom computational pipeline, and barcodes were selected as protospacers based on specific criteria (**Fig. S9**). To normalize the initial abundance of selected targets, we intentionally chose ultra-rare barcodes with only 1–2 reads in the original libraries, due to large variation in read counts per barcode. In total, 1, 384 sgRNAs were designed to target 684 perfect DHFR genes in the 1, 536-gene library, and 389 sgRNAs were designed to target 149 genes in the 384-gene library.

The 1, 384-plex and 389-plex sgRNA libraries were *in vitro* transcribed from a microarray-derived oligo pool (46). However, RNA-seq revealed severe spacer bias in these libraries, likely stemming from T7 RNAP preference for templates with a 5′ guanine tetramer (5’ GGGG) immediately downstream of the T7 promoter (46, 71). As the initial sgRNA libraries were synthesized with only a single 5′ G to initiate transcription, spacer representation was highly unequal, though these libraries were still used for large-scale enrichment experiments.

To address this bias, we explored adding a 5′ GGGG sequence to all spacer templates and compared the resulting uniformity against the 5’ G condition. While the sgRNA libraries used here predated full optimization, we sought to determine whether the 5′ GGGG modification impacted log₂ enrichment.

For this comparison, we subdivided the 389-plex library into smaller groups of 12, 60, and 389 spacers, targeting an equivalent number of barcodes corresponding to perfect assemblies from the 384-gene library. All templates were padded with a 5′ GGGG except a duplicate of the 12-spacer subpool, which retained a single 5′ G to directly compare the two conditions. The sgRNA libraries were *in vitro* transcribed in emulsions (46) and subsequently used for BAR-CAT enrichment.

To assess the impact of target scale on BAR-CAT performance, we enriched 12, 60, and 389 barcodes from the 384-gene DHFR plasmid library, and 1, 384 barcodes from the 1, 536-gene plasmid library, using sgRNAs containing either a 5′ G or 5′ GGGG. A synthetic sgRNA targeting a single barcode (Integrated DNA Technologies) served as a singleplex enrichment control.

A small-scale enrichment performance baseline was established from singleplex (n = 2) and 12-plex (5′ G, n = 2; 5′ GGGG, n = 1) enrichments. Two replicates of single-barcode enrichment achieved 8.4-fold and 6.8-fold log₂ enrichment, corresponding to 349-fold and 115-fold linear values, respectively. Enrichment of 12 barcodes with 5′ G sgRNAs resulted in a 6.9-fold median log₂ enrichment (122-fold) (**Fig. 3A**) and 54-fold population fraction enrichment with zero target dropouts. A second replicate yielded a 3.5-fold median log₂ enrichment (11-fold) with a 16-fold population fraction enrichment and 18.2% target dropout (**Fig. 3B**). Enrichment using a 5′ GGGG sgRNA library produced similar results (log₂ enrichment of 4.8 [29-fold] and 18.2% target dropout).

**Fig. 3.**
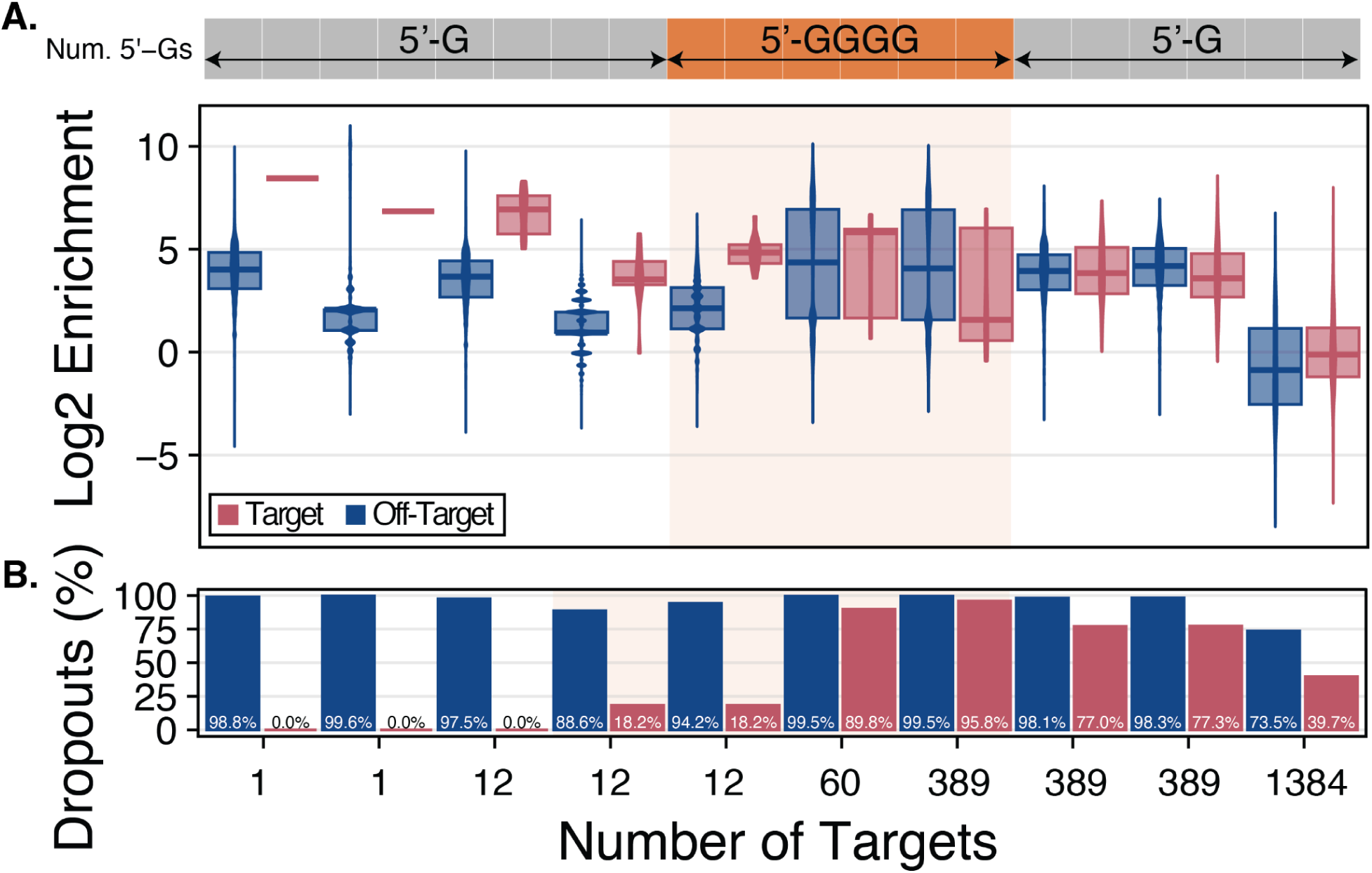
BAR-CAT v1.0 performance declines with increasing enrichment scale and is further reduced by sgRNA spacers starting with 5′ guanine tetramers (5′ GGGG) compared to single 5′ guanine (5′ G) designs. **A.** Overlaid violin and boxplots show log₂ enrichment scores (log₂ fold change in barcode abundance before vs. after enrichment) for target (magenta) and non-target (blue) barcodes across different enrichment scales and sgRNA 5′ guanine modifications. Shaded areas indicate the interquartile range; bars mark the median. The colored bar above each distribution indicates sgRNA design: 5′ G (gray) or 5′ GGGG (orange). Singleplex enrichments (5′ G) from the 384-gene DHFR library (library S4) achieved median log₂ enrichments of 6.8 and 8.4, corresponding to 115- and 349-fold enrichment. At the 12-plex scale, median values dropped to 3.5 and 6.9 (11.6- and 122-fold), with 16.3-and 349-fold population fraction enrichment using 5′ G sgRNAs. For 5′ GGGG sgRNAs, the median enrichment was 4.8 (29-fold), with 26-fold population fraction enrichment. At 60-plex, the median log₂ enrichment was 5.8 (57-fold), with 3.7-fold population fraction enrichment. At the 389-plex scale, 5′ GGGG sgRNAs yielded a median log₂ enrichment of 1.6 (3.0-fold) with 0.7-fold population fraction enrichment, while 5′ G sgRNAs performed modestly better (3.5 and 3.8, or 12.0- and 14.3-fold, with 5.0-fold population fraction enrichment). Enrichment at 1, 384-plex scale from the 1, 536-gene DHFR library (library S2) was negligible. These reductions likely stem from sgRNA competition for dCas9 at larger scales (72, 73), and, for 5′ GGGG sgRNAs, dilution of active complexes by excess high molecular weight (HMW) byproducts (46). **B.** Percent dropout values are listed in the bar plots corresponding to off-target (blue) and target (magenta) barcodes according to the scale of targeted barcodes, as indicated by the x-axis. For 12-plex enrichments, target dropouts were 0.0% (5′ G) and 8.3% (5′ G and 5′ GGGG). Dropouts increased to 89.8% at 60-plex (5′ GGGG), 95.8% for 389-plex (5′ GGGG), and 77.0% and 77.3% for 389-plex (5′ G). In contrast, the 1, 384-plex enrichment showed 39.7% target dropouts, likely reflecting improved barcode diversity in the 1, 536-gene DHFR plasmid library. Off-target dropouts remained high across all scales (73.5–99.6%).

However, sgRNA age appeared to impact performance, as indicated by a 3.4-point decrease in median log₂ enrichment for the 12-plex sgRNA library after just 10 weeks. In contrast, the synthetic single sgRNA used for DHFR enrichment showed only a 1.6-point decrease in median log₂ enrichment over 20 weeks. The 3-plex synthetic sgRNAs showed virtually no change during the first 20 weeks, followed by a 5.1-point drop in the final 20 weeks (**Fig. S8**). These results suggest that sgRNA degradation over time likely contributed to reduced enrichment efficiency. Nonetheless, in the 12-plex context, sgRNAs with a 5′ GGGG sequence did not exhibit lower enrichment values than those with a single 5′ G.

We next examined off-target barcode dropouts. Singleplex enrichments had 98.8% and 99.6% off-target dropouts, while 12-plex enrichments showed 97.5% (5′ G), 88.6% (5′ G), and 94.2% (5′ GGGG) (**Fig. 3B**). These values were markedly higher than those observed in all *rfp* library enrichments, which had a mean off-target dropout of 5.73% ± 2.52% (range: 1.5–10.2%) (**Fig. 2B**, **Fig. 1D**), corresponding to 91.8% (5′ G), 82.9% (5′ G), and 88.5% (5′ GGGG) increases, respectively. Since bead washing conditions were unchanged between BAR-CAT v0.3 and v1.0, we attributed the elevated off-target dropout to the much larger barcode population in the 384-gene library (5, 300, 938 barcodes) compared to the *rfp* library (306, 388 barcodes). The greater barcode diversity likely increased the probability of off-target loss during washing.

Fractional barcode distribution analysis revealed that low-abundance off-target barcodes were consistently enriched from approximately 10^-7^–10^-6^ to 10^-4^–10^-3^ across both singleplex and 12-plex enrichments regardless of spacer design. This off-target enrichment likely masked the enrichment of true targets, especially at small scales (**Fig. S10A-E**). Moreover, since only 5 out of 5, 300, 938 barcodes had abundances above 10^-5^, most high-abundance off-target barcodes likely dropped out, while low-abundance barcodes had a higher chance of enrichment by chance alone.

Next, we evaluated whether the trends observed for small-scale enrichment from the 384-gene DHFR library extended to larger-scale enrichment targeting 60 (5′ GGGG, n = 1), 389 (5′ G, n = 2), and 389 (5′ GGGG, n = 1) barcodes. The 60-plex enrichment yielded a 5.8-fold median log₂ enrichment (57-fold) (**Fig. 3A**) with a 3.7-fold population fraction enrichment and with 89.8% of target barcodes dropping out (**Fig. 3B**). In contrast, the 389-plex enrichment with 5′ GGGG-modified sgRNAs resulted in a 1.6-fold median log₂ enrichment (3.0-fold) (**Fig. 3A**), a negligible 0.7-fold population fraction enrichment, and 95.8% target dropouts (**Fig. 3B**). The poor performance of sgRNAs with the 5′ GGGG modification at larger scales suggests this modification may hinder efficiency compared to 5′ G sgRNAs at higher multiplexing scales.

In contrast, 389-plex enrichments using 5′ G-modified sgRNAs achieved median log₂ enrichments of 3.8 (14-fold) and 3.6 (12-fold), representing 4.7- and 4.0-fold increases compared to the same condition with 5′ GGGG-modified sgRNAs (**Fig. 3A**). These enrichments yielded a 5.0-fold population enrichment and a mean target dropout rate of 77.2% ± 0.21 (n = 2), compared to 95.8% in the 5′ GGGG condition (n = 1), suggesting an 18.6% reduction in dropout (**Fig. 3B**). Together, these results indicate that the poor performance at large scales is intrinsic to BAR-CAT v0.1 and is exacerbated by sgRNA design, with 5′ GGGG-modified sgRNAs performing substantially worse than 5′ G-modified sgRNAs.

We attribute the inferior performance of 5′ GGGG-modified sgRNAs to the production of excessive high molecular weight (HMW) RNA species during IVT. Because the same mass of sgRNA was added to each enrichment condition, the presence of high molecular weight (HMW), inactive RNAs diluted the effective concentration of functional sgRNA spacer molecules, disproportionately reducing enrichment efficiency at higher multiplexing levels (46). Based on these findings, we recommend avoiding 5′ GGGG sgRNAs for large-scale applications. Instead, sgRNAs should be transcribed with a single 5′ G preceding the spacer sequence either in emulsions or using high template input (e.g., 400 ng DNA in 20 μL IVTs), both of which improve sgRNA uniformity without introducing excessive HMW RNA species (46).

We next examined the distribution of target and off-target barcode fractions before and after enrichment for the 60-plex and 389-plex DHFR library enrichments. Consistent with previous results (**Fig. S10A-E**), off-target barcodes initially present at 10^-7^–10^-6^ were enriched to 10^-4^–10^-3^ following enrichment (**Fig. S11A-E**). For both the 60-plex (**Fig. S11A**) and 389-plex (**Fig. S11C**) conditions, target and off-target barcode distributions overlapped after enrichment, with only a small subset of target barcodes enriched above background. Most target barcodes dropped out, while enrichment of low-abundance off-targets obscured the enrichment signal. High-abundance off-target barcodes from the original library were largely depleted. A similar pattern was observed for the two 389-plex (5′ G) replicates (**Fig. S11D, E**), although slightly more target barcodes were recovered compared to 5′ GGGG conditions, consistent with reduced dropout rates.

These observations suggest that the high barcode diversity of the 384-gene DHFR plasmid library led to enrichment of numerous low-abundance off-target barcodes, introducing substantial noise into the results.

Applying a more stringent bottleneck to reduce overall library diversity should therefore mitigate this issue and improve enrichment performance. This prediction is supported by our finding that off-target enrichment patterns were primarily driven by library composition, although sgRNA design (5′ GGGG vs 5′ G) also influenced target dropouts.

With this in mind, we assessed the performance of the largest enrichment condition tested: enrichment of 1, 384 targets from the 1, 536-gene DHFR plasmid library (library S2) using 5′ G sgRNAs (n=1). This condition yielded a negative 0.11-fold median log₂ enrichment (0.92-fold) (**Fig. 3A**) with a population fraction enrichment of 1.2-fold and 39.8% target dropouts (**Fig. 3B**), indicating no meaningful targeted enrichment. Although enrichment was negligible at the 1, 384-plex scale compared to 389-plex, the percentage of target dropouts was lower compared to the 389-plex enrichments (39.8% vs. a mean of 77.2% ± 0.21 (n = 2) for 5′ G and 95.8% (n = 1) for 5′ GGGG). The population fraction of off-target barcodes did not show enrichment (1.0-fold) for the 1, 384-plex condition, similarly to the 389-plex conditions. Additionally, the 1, 384-plex condition had 73.5% off-target dropouts, about 23.6% less compared to the mean of 97.1% ± 3.64% (n = 9) obtained across all enrichments prepared with the 384-gene DHFR plasmid library regardless of scale (**Fig. 3B**).

These results suggest that, although the 1, 384-plex enrichment did not yield strong overall enrichment, it showed reduced dropout of both target and off-target barcodes. We hypothesize that this improvement is due to lower diversity of the 1, 536-gene DHFR plasmid library (S2), in contrast to the 384-gene DHFR plasmid library (S4), which contained a much higher proportion of high-abundance target and off-target barcodes. Supporting this, a comparison of spacer fractions in the enriched versus original populations revealed that most targeted barcodes in the 1, 384-plex condition were retained and modestly enriched as a distinct subpopulation, while off-target barcodes were also retained. Both sets were largely clustered around the unity line (**Fig. S11B**), indicating minimal change in representation. Altogether, these observations support our hypothesis that high barcode diversity in the smaller DHFR plasmid library contributed to excessive dropout of both target and high-abundance off-target barcodes.

However, although the lower barcode diversity in the 1, 536-gene DHFR library (S2) likely contributed to better retention of target barcodes in the 1, 384-plex enrichment, it does not fully explain the absence of meaningful targeted enrichment under this condition. We hypothesized that the low enrichment resulted in part from increased competition among the larger number of sgRNA spacers for free dCas9, which limited sgRNA-mediated enrichment for most targets. Nevertheless, since some barcodes still achieved strong enrichment while most did not (**Fig. S11B**), we reasoned that additional factors might differentiate high-performing from low-performing targets. Therefore, we took a closer look at the features of individual barcodes to better understand the underlying causes of variable performance.

### Evaluating sgRNA Performance, sgRNA Library Bias, and Barcode Target Selection to Improve Future Large-Scale BAR-CAT Studies

We examined whether the log₂ enrichment of targeted barcodes in the 389-plex and 1, 384-plex experiments correlated with predicted sgRNA on-target efficiencies to investigate the lack of targeted enrichment (**Fig. 3A**) and the variability in sgRNA spacer performance (**Fig. S11B**). We used CRISPRscan, which predicts sgRNA performance based on *in vivo* editing efficiencies measured in zebrafish embryos (74). However, comparing these predicted CRISPR efficiency scores to the observed log₂ enrichment values from BAR-CAT v0.1 (excluding sgRNAs with dropped-out targets) showed no significant correlation for either the 389-plex enrichments from the 384-gene DHFR library (5′ G sgRNAs, replicate 2, R² = 0.009; **Fig. S12A**) or the 1, 384-plex enrichments from the 1, 536-gene DHFR library (5′ G sgRNAs, R² = 0.015; **Fig. S12B**). This lack of correlation was not unexpected, as most prediction tools are developed for *in vivo* genome editing rather than *in vitro* systems like BAR-CAT. Similarly, Henser-Brownhill and colleagues observed minimal correlation between sgRNA activity and predicted scores using sgRNA Scorer 2.0 (75, 76), consistent with other studies that have highlighted the limitations of predictive algorithms for *in vitro* applications (77, 78). While this mismatch between predictions and our assay provides one possible explanation, other factors beyond sgRNA efficiency may be affecting performance in our system.

Next, we explored whether the distribution of spacers within the sgRNA libraries, as determined by RNA-seq in our prior work (46), influenced the log₂ enrichment of targeted barcodes in the 389-plex and 1, 384-plex experiments. We hypothesized that uneven spacer abundance following *in vitro* transcription of sgRNA libraries could hinder targeted enrichment with BAR-CAT at larger scales. However, no correlation was found between spacer distribution and observed log₂ enrichment for either the 389-plex (5′ G sgRNAs, replicate 2; **Fig. S13A**) or the 1, 384-plex (5′ G sgRNAs; **Fig. S13B**) experiments. This lack of correlation suggests that, despite substantial variation in sgRNA spacer distribution, other factors such as dCas9 binding dynamics or local sequence context may play a more dominant role in determining enrichment efficiency.

Additionally, as the number of sgRNAs increases, they compete for free dCas9, reducing the overall activity of all sgRNAs (72, 73). This phenomenon was highlighted in the development of a negative feedback circuit designed to increase dCas9 expression as free dCas9 becomes saturated due to co-expression of multiple sgRNAs (79). The impact of sgRNA competition is particularly dramatic. For instance, one study using dCas9 as a repressor reported that repression decreased from 58-fold with a single sgRNA to 10-fold when seven sgRNAs were co-expressed (72). Furthermore, another study showed that dCas9 saturation occurs when approximately 12 sgRNAs compete for binding (80), which may explain the high target dropouts observed during enrichment of more than 12 targets from the DHFR libraries. As a result, we predicted that increasing sgRNA, dCas9, or DNA inputs could mitigate this effect and improve enrichment scores. More specifically, we wanted to increase the DNA input amount to mitigate any increased off-target effects from increasing sgRNAs or dCas9, since it reduces the RNP to DNA ratio, as described previously (**Figs. 1D, 2A**).

To test this, we enriched 12 targeted barcodes from the 384-gene DHFR plasmid library (library S4) while varying the sgRNA, dCas9, or DNA inputs to evaluate their impact on enrichment. We performed the standard BAR-CAT v1.0 protocol using 12-plex sgRNA pools with spacers beginning with either a 5′ G or 5′ GGGG. Across all conditions, the percentage of target barcode dropouts was high (25–66.7%) compared to those previously observed for the 12-plex library (0% and 18.2% target dropouts; **Fig. 3B**), likely due to issues with dCas9 reconstitution. Nonetheless, the results provided general preliminary trends. The 5′ G control showed a log₂ median enrichment of 4.6 (24-fold) and a 16-fold population fraction enrichment with 36.4% target dropouts. The 5′ GGGG control yielded a log₂ median enrichment of 6.3 (77-fold) and a 29-fold population fraction enrichment with an even higher percentage of target dropouts (63.6%). An experimental condition with 3× more input sgRNAs (5′ G) resulted in a log₂ median enrichment of 3.9 (14-fold) and 9.1-fold population fraction enrichment, markedly lower than the corresponding 5′ GGGG or 5′ G controls (**Fig. S14**). There were 27.3% target dropouts, lower than either control condition. Simultaneously increasing both DNA (2.2×) and dCas9 (3×) led to a median log₂ enrichment of 4.0 (16-fold) and 14-fold population fraction enrichment with 18.2% target dropouts. An experimental condition with 2.2× DNA and 3× sgRNAs produced a median log₂ enrichment of 3.1 (8.6-fold) and 4.1-fold population fraction enrichment with 54.5% target dropouts. Lastly, we tested a 2× reaction volume condition in which sgRNAs were scaled by 2×, DNA by 5×, and dCas9 by 3×. This condition yielded a median log₂ enrichment of 4.3 (20-fold) and a 13-fold population fraction enrichment with 36.4% target dropouts **(Fig. S14**).

Overall, these results showed that increasing sgRNA or dCas9 concentrations did not substantially improve enrichment at the 12-plex scale, and increasing input library DNA did not rescue this effect. In fact, enrichment scores often decreased, potentially due to greater off-target barcode enrichment due to excess active RNPs. Off-target barcode dropout rates remained consistent across conditions, with 5′ GGGG and 5′ G controls showing 48.0% and 41.0% dropout, respectively, and a mean of 43.7% ± 2.61 (**Fig. S14**), indicating that the tested variables had minimal impact on this metric. Comparing enriched versus original barcode fractions across all conditions did not differ from those previously observed for 12-plex enrichments with the 384-gene DHFR plasmid library (library S4) (**Fig. S15A–F**); both low-abundance off-targets and target barcodes that did not drop out were enriched. These results indicate that varying DNA, sgRNA, or dCas9 input has little effect on enrichment efficiency at small scales. However, we anticipate that increasing dCas9 concentration will have a greater impact at larger scales (e.g., 389+ targets), where competition among sgRNAs for dCas9 is more pronounced. Thus, increasing dCas9 levels during RNP assembly, while keeping sgRNA levels constant, may reduce competition and ensure sufficient complex formation per target.

Because enrichment of target barcodes from the DHFR gene libraries was accompanied by off-target enrichment, we sought to better characterize the identities of the enriched off-target barcodes. We first examined whether enriched off-target barcodes contained perfect matches to the 7–10 base pair PAM-adjacent seed region of any of the 18 target spacers. However, we found no significant enrichment of off-target sequences with seed matches compared to those without seed matches (data not shown). This finding is surprising, as previous studies have shown that CRISPR off-target effects are primarily driven by seed sequence matches (41, 42, 55, 69).

One possible explanation for this lack of correlation is the promiscuous binding behavior of dCas9. Both dCas9 and Cas9 can bind off-target sites with a PAM, even when mismatches occur in the seed region, and this binding may persist unless mismatches are also present in the RDR (56). Although mismatches in the seed region generally reduce binding efficiency (42, 81), off-target enrichment could result from numerous partial sequence matches across a wide range of sequences in complex libraries like the DHFR libraries (82). While we cannot definitively explain why seed sequence similarity did not predict off-target enrichment in this case, we propose that dCas9’s binding promiscuity, in combination with other experimental variables, contributes to the observed off-target enrichment trends. This suggests that factors such as the specific sequence context, spacer distribution, or RNP concentration may also influence off-target behavior in high-complexity libraries.

Since off-target enrichment does not appear to be driven by seed sequence similarity, re-designing target sequences may not significantly reduce off-target noise. Instead, we propose targeting barcodes based on their original abundance as a more effective strategy. For example, the three barcodes selected for BAR-CAT optimization (**Fig. 2**) were chosen from a set of 18 barcodes (**Fig. 1B**) spanning a range of abundance. Barcodes 14 and 15, selected from the low-to-medium abundance range, exhibited the best log₂ enrichment scores following BAR-CAT v0.3 (proteinase K denaturation, **Fig. 1D. Fig. S6**). In contrast, enrichment efficiency for both the most abundant (BCs 1–13) and least abundant (BCs 16–18) barcodes declined (**Fig. S6**). This is because it is inherently easier for low abundance targets to achieve high fold enrichments. When fold change is normalized to the starting copy number, a barcode present at a single read needs only nine additional captures to register a 10-fold gain, whereas one that begins at 1, 000 reads must accumulate 9, 000 more molecules, a requirement that rapidly encounters binding and amplification saturation limits. In other words, high-abundance targets may show lower fold-enrichment because further amplification has limited impact on already abundant sequences, while targeting ultra-low-abundance barcodes could increase off-target enrichment as RNPs search for rare targets. Therefore, targeting barcodes with low-to-medium abundance may provide a "sweet spot" for efficient enrichment. The robust enrichment with reduced noise observed in **Fig. 2** for barcodes 7, 8, and 15 may stem from this strategy. However, this approach may not be effective in very diverse libraries, like the 384-gene DHFR library (S4), where barcode distributions are skewed towards low abundance, leading to frequent target dropouts and higher off-target binding.

An alternative approach to improving both accuracy and scalability involves transitioning from dCas9 to catalytically active Cas9 in a modified version of BAR-CAT. Cas9 could enable more stringent target selection by cleaving DNA, unlike dCas9, which only binds to DNA. The ability of Cas9 to cleave even ultra-low abundance barcodes would facilitate their amplification by PCR, whereas dCas9-based methods rely on higher target abundance for effective enrichment. A Cas9-based strategy could linearize supercoiled targets within the gene library via cleavage. This would enable adapter ligation and the generation of unique PCR handles for selective amplification, similar to the FLASH method, which enriches low-abundance pathogenic DNA from patient samples (77). Cas12a-Capture offers a similar approach to FLASH, using Cas12a to cleave genomic targets associated with Joubert Syndrome, allowing efficient adapter ligation and targeted sequencing while avoiding the limitations of dCas9-based methods (83). Compared to dCas9 enrichment, which relies on bead washing and often retains off-targets, these cleavage-based methods may provide cleaner enrichment. While a Cas9-based BAR-CAT would necessitate additional steps such as cloning and ligation post-enrichment, it could also expand BAR-CAT’s utility to targeted sequencing of rare alleles or pathogenic variants. However, if Cas9 exhibits non-specific cleavage, as reported in the literature (45), its off-target effects may be comparable to those observed with dCas9.

There are also distinct advantages to using dCas9 for BAR-CAT, particularly in applications involving cloning and length-independent DNA enrichment. First, increasing the number of targets enriched with our dCas9-based BAR-CAT approach could simplify direct cloning of supercoiled perfect genes into *E. coli*, making downstream screening and functional validation more efficient than with a Cas9-based system. Second, a key benefit of dCas9 is that it does not cleave DNA, making BAR-CAT enrichment inherently length-independent in principle. In BAR-CAT v1.0, barcodes were successfully targeted from ∼2.7 kbp supercoiled gene libraries, and the corresponding plasmids were retrieved in full, demonstrating the method’s capacity to enrich DNA of this length.

In practice, recovery of longer DNA fragments can be hindered by several factors. Early studies indicated that mechanical shear stress, such as pipetting, often fragmented DNA into heterogeneous distributions of 0.5–10 kbp (84). However, modern protocols, including slow-pipetting techniques to reduce shearing of 100 kbp DNA (85), have improved the handling of HMW DNA, making pipetting less of a limiting factor. We predict that BAR-CAT may show reduced recovery of DNA targets >20 kbp due to slower diffusion kinetics and increased steric hindrance during bead capture of dCas9 and subsequent washing steps. These factors suggest that while enrichment of fragments up to 10 kbp is feasible, recovery efficiency may decline with increasing fragment size, depending on fragment design and reaction conditions. Direct experimentation will be required to accurately assess the DNA length limitations of BAR-CAT. Based on current predictions, BAR-CAT should be effective for enriching sequences up to 20 kbp with minimal issues.

In addition to synthetic gene libraries, dCas9-based BAR-CAT could theoretically be applied to enrich ancient DNA for targeted next-generation sequencing. This approach offers a potential alternative to RNA hybridization-based methods (86), which may be limited by mismatches or degradation. In this context, dCas9 could bind conserved housekeeping genes without cleaving the fragile, partially degraded DNA, enabling recovery of fragments even when only a few conserved targets are available. Thus, a highly multiplexed and optimized version of BAR-CAT could serve as a valuable tool for enriching both synthetic genes and ancient DNA fragments up to ∼20 kbp in length, leveraging the inherent length-independence of dCas9 binding and pull-down.

Currently, BAR-CAT v1.0 successfully enriches 3–12 unique targets simultaneously, demonstrating its potential and providing valuable insights for further method development. However, improving its scalability is critical to fully achieving the original goals of BAR-CAT. To address this, we plan to titrate dCas9 concentrations during enrichment reactions to better manage sgRNA competition as library complexity increases. Additionally, we aim to selectively target barcodes with low to moderate abundance in DropSynth libraries to minimize off-target enrichment. We will also investigate whether sequential enrichment can boost the representation of targeted barcodes. For this, we hypothesize that targeting low-abundance barcodes corresponding to perfect gene assemblies will result in moderate enrichment levels. These moderate levels could then be enhanced by repeating the BAR-CAT process prior to final amplification, effectively driving moderately enriched barcodes to higher abundance and reducing off-target enrichment and noise. Finally, we will evaluate a modified version of BAR-CAT using Cas9 instead of dCas9 to assess whether direct cleavage can improve both enrichment specificity and scalability by enabling selective amplification of cleaved targets.

## Conclusions

Although the application of plasmid DNA libraries has evolved from gene discovery and early sequencing efforts to large-scale vehicles of functional discovery, their role as drivers of molecular biology has not diminished. However, as massively parallel assays, protein engineering, functional genomics, and synthetic biology applications continue to expand, so will the demand for larger DNA libraries with longer genes. DropSynth is one notable method that aims to meet this demand, but as other assembly methods, it suffers from the significant accumulation of oligo errors at long lengths >1 kbp. To address this issue, we developed BAR-CAT, a method that applies CRISPR-dCas9 to selectively enrich perfect gene assemblies.

During its proof-of-concept, BAR-CAT enriched 18 unique barcodes distributed from high to low abundance within a barcoded *rfp* library by a median of 6.3-fold. Increasing the cumulative wash volume of the magnetic beads from 0.3 mL (6 × 50 µL) to 18 mL (9 × 2 mL) boosted the median enrichment to about 1.9-fold. Additionally, non-enriched off-targets were further depleted following enrichment, reducing background noise.. Meanwhile, adding a dCas9 denaturation step to minimize non-specific interactions between dCas9, DNA, and beads did not increase enrichment but contributed to increased off-target barcode dropout. Lastly, increasing the input DNA from 50 ng to 500 ng led to a median 600-fold enrichment value and an impressive 1, 094-fold population fraction enrichment for three low-to-median abundance barcode targets in the *rfp* library. These improvements likely stem from adjusting the RNP-to-DNA ratio by adding more DNA, thereby limiting excess RNPs that could otherwise bind off-targets. All of these process optimizations were incorporated into BAR-CAT version 1.0, as presented in this work.

When BAR-CAT was applied to enrich ultra low-abundance targets from 384- and 1, 536-member DHFR plasmid libraries assembled by DropSynth, we observed excessive target dropouts exceeding 90% in some cases. At the 389-plex scale, some of these effects were attributable to sgRNA libraries containing a 5′ GGGG sequence, which resulted in insufficient sgRNA concentrations and excessive HMW products. However, we believe that additional factors, including very high diversity of the DHFR library, targeting of extremely low-abundance members, and sgRNA competition for dCas9, contributed to robust off-target effects and severe dropout of targets. Future work will address these shortcomings and explore alternatives such as using Cas9 in place of dCas9 and implementing sequential enrichment strategies.

In conclusion, the development of BAR-CAT v1.0 provides a practical framework for researchers aiming to design CRISPR-based DNA enrichment methods. It offers insight into the nuances of enrichment behavior in a simplified yet still complex system. We hope that the potential utility of BAR-CAT, once further optimized, will inspire the development of additional tools for applications such as synthetic gene enrichment, ancient DNA enrichment, diagnostics, and targeted next-generation sequencing.

## Supporting information

Experimental Data Summary

## Acknowledgements

This work was supported by the National Science Foundation MCB-2032259 grant. Research reported in this publication was supported by the National Institute of General Medical Sciences of the National Institutes of Health under award number T32GM149387 (to NV). The content is solely the responsibility of the authors and does not necessarily represent the official views of the National Institutes of Health. Special thanks to Yuki R. Gaudreault for assisting with some of the experimental work.

## Data and Materials Availability

Raw Illumina MiSeq reads of the target gene libraries and nanopore sequence reads for enriched libraries were submitted to the NCBI Sequence Read Archive under BioProject accession PRJNA1273454 (https://www.ncbi.nlm.nih.gov/bioproject/1273454). Processed enrichment data for each spacer for all experiments is available on FigShare (https://doi.org/10.6084/m9.figshare.29428988)

## Methods

### Constructing the pEVBC3 barcoding plasmid for the rfp library

Plesa and colleagues developed pEVBC1, a pUC19-derived plasmid that adds 20-mer barcodes to cloned genes and expresses them in *E. coli* via a pLac-UV5 promoter (15). Since pEVBC1 was available in-house, a new plasmid, designated pEVBC3, was derived from pEVBC1 to develop a pilot gene library using PCR-round-the-horn. Four alternative PAM sites, an EcoRI restriction site, and the pLac-UV5 promoter were removed.

To prepare the vector backbone, the parent plasmid pEVBC1 was linearized by double digestion with XbaI and KpnI-HF (New England Biolabs, Ipswich, MA, USA), which removed the *mCherry* cargo gene. Each 50 µL reaction contained 500 ng of pEVBC1, 5 µL of 10X CutSmart Buffer (NEB), 1.0 µLof XbaI (20, 000 units per milliliter), 1.0 µL of KpnI-HF (20, 000 units per mL), and nuclease-free water. The digestion was incubated at 37 °C for one hour and heat inactivated at 65 °C for 20 minutes. The linearized plasmid was purified using DNA Clean and Concentrator columns (5 µg capacity, Zymo Research Corp, Irvine, CA, USA). A 1, 869 bp product corresponding to the digested vector was verified by 1% agarose gel electrophoresis alongside a 1 kbp DNA ladder (New England Biolabs).

To build pEVBC3, A series of three consecutive PCR protocols was performed to complete PCR round-the-horn (**Table S3**), following the method previously described in prior work (15). The first PCR aimed to modify the linearized pEVBC1 into pEVBC3. Eight PCR reactions were set up, each containing 0.5 ng of linearized pEVBC1, 0.5 µL each of 10 µM forward (EVBC3_FWD1) and reverse (EVBC3_REV1) primers (**Table S4**), 25 µL of 2X Q5 High-Fidelity Master Mix (NEB), and nuclease-free water to a total volume of 50 µL. The thermocycler PCR program included an initial denaturation at 98 °C for 30 seconds, followed by 35 cycles of 98 °C for 10 seconds, annealing at 72 °C for 30 seconds, extension at 72 °C for 1 minute per cycle, and a final extension at 72 °C for 2 minutes. The resulting 1, 950 bp pEVBC3 backbone (PCR 1 product) was purified using 5 µg DNA Cleanup Columns (Zymo Research Corp, Irvine, CA).

The purpose of the second PCR was to add a randomly-generated 20-mer sequence to barcode genes cloned into pEVBC3. Each individual PCR was prepared with 1 ng of PCR 1 product, 2.5 µL each of 10 µM forward (EVBC3_FWD2), and reverse (EVBC3_REV2) primers (**Table S4**), 25 µL of 2X Q5 High-Fidelity Master Mix (NEB), and nuclease-free water to a final volume of 50 µL. The same thermocycler PCR program was used as in PCR 1, except that only five amplification cycles were performed. The resulting 2 kbp barcoded pEVBC3 backbone (PCR 2 product) was purified using 5 µg DNA Cleanup Columns (Zymo Research Corp, Irvine, CA).

The purpose of the third PCR was to bulk amplify the barcoded pEVBC3 backbone in preparation for cloning in gene libraries. Each individual PCR was prepared with 1 µL of PCR 2 product, 2.5 µL each of 10 µM forward (EVBC3_FWD3RE), and reverse (EVBC3_REV2) primers (**Table S4**), 25 µL of 2X Q5 High-Fidelity Master Mix (NEB), and nuclease-free water to a total volume of 50 µL. The same thermocycler PCR program was used as in PCR 1, except that only 15 amplification cycles were performed. The amplified pEVBC3 backbone (PCR 2 product) and 993 bp PCR 3 product were verified with a 1% agarose gel and 1 kbp ladder (New England Biolabs, NEB, Ipswich, USA). The pEVBC3 backbone (2.7 kbp) was verified via Sanger Sequencing following ligation to an *mCherry* gene, transformation into NEB® 5-alpha Competent *E. coli*, and colony picking.

### Amplification of mCherry red fluorescent protein (rfp) gene for pilot library generation

Bulk amplification of a 708 bp *mCherry* gene was performed to obtain sufficient insert material for generating a single-gene *rfp* library. The library was later barcoded using pEVBC3. The original *mCherry* sequence was sourced from pZS2-123, a plasmid gifted to Dr. Sri Kosuri from Michael Elowitz (Addgene plasmid #26598; http://n2t.net/addgene:26598; RRID: Addgene_26598). The Kosuri group derived pSK48, the plasmid containing amplified *mCherry* used in this work, from its parent plasmid pZS2-123.

In total, 30 PCRs were performed to obtain 5-10 µg of bulk amplified, biotinylated *mCherry* for downstream cloning. Each reaction contained 1.0 ng of pSK48, 2.5 µL each of 10 µM forward (RFP_KpnI_FWD_Biotin_NV) and reverse (RFP_Ndel_REV_Biotin_NV) primers (**Table S2**), 25 µL of 2X Q5 High-Fidelity Master Mix (NEB), and nuclease-free water to a total volume of 50 µL. The thermocycler PCR program included an initial denaturation at 98 °C for 30 seconds, followed by 35 cycles of 98 °C for 10 seconds, annealing at 72 °C for 30 seconds, extension at 72 °C for 30 minute per cycle, and a final extension at 72 °C for 2 minutes. The *mCherry* PCR product was purified using 5 µg DNA Cleanup Columns (Zymo Research Corp, Irvine, CA).

### Constructing the pEVBC8 barcoding plasmid for DropSynth dihydrofolate reductase (DHFR) libraries

The main modification made to the pEVBC3 barcoding plasmid was the addition of a protospacer adjacent motif (PAM) site to the 3′ end of the barcode region. To prepare the vector backbone, pEVBC3 containing an *mCherry* insert (pEVBC3-RFP) and the barcode TAAATATTACCGGTCTCTTT was purified using 5 µg DNA Cleanup Columns (Zymo Research Corp, Irvine, CA). The sequence of this plasmid was verified by Sanger sequencing and then linearized with KpnI-HF® (New England Biolabs, Ipswich, USA) and NdeI (NEB) restriction enzymes. Each digestion included 666 ng of pEVBC3, 5 µL of 10X CutSmart® Buffer (NEB), 1.0 µl of KpnI-HF® (20, 000 units/mL) (NEB), 1.0 µL of NdeI (20, 000 units/mL) (NEB), and nuclease-free water to a total volume of 50 µL. The digest was incubated in a thermocycler at 37 °C for 1.5 hours and heat inactivated at 65 °C for 20 min. The products were purified using 5 µg DNA Cleanup Columns (New England Biolabs, Ipswich, USA). The 2 kbp pEVBC3 backbone was size-selected from a 1.0% agarose gel using the Monarch® DNA Gel Extraction Kit (NEB), with a 1 kbp ladder (NEB) for comparison.

To build pEVBC8, we used the same PCR-round-the-horn protocol for producing pEVBC3 but with a few modifications (**Table S3**). The first PCR included 1 ng of linearized pEVBC3, a forward primer (EVBC8_FWD1), and 0.625 µL of each 10 µM primer (6.25 µM final concentration). The thermocycler program used was identical to that used in the first PCR for generating pEVBC3. The 1, 950 bp PCR 1 product was size-selected from a 1.0% agarose gel using the Monarch® DNA Gel Extraction Kit (NEB). Modifications to the second PCR included the addition of a forward primer (EVBC8_FWD2; **Table S4**) to eliminate a potential alternative PAM site on the 5’ end of the barcode. Additionally, 0.625 µL of each 10 µM primer (6.25 µM final concentration) was added. The third PCR was performed as described for generating pEVBC3, however, 60 PCRs were prepared and 1 ng of PCR 2 product was added instead of 1 µL. The final amplified pEVBC8 backbone was validated by agarose gel electrophoresis (1.0% agarose gel) and nanopore sequencing (Plasmidsaurus).

### DropSynth assembly of DHFR gene libraries and amplification

Four DHFR DropSynth libraries were assembled and amplified to obtain gene library insert material, later barcoded using pEVBC8. Library S4 consisted of 384 genes, ranging from 473 to 545 bp in length, assembled from Twist Bioscience 300-mer oligos. In contrast, library S2 contained 1, 536 genes, ranging from 464 to 548 bp in length, assembled from HiFi SurePrint High Fidelity (HiFi) 230-mer oligos (Agilent, Santa Clara, USA). Both libraries were synthesized using DropSynth 2.0, with library S2 assembled by a prior study (17) and library S4 assembled for this study.

Prior to bulk amplification of the 384-gene DHFR library (S4), qPCR (CFX Opus 96, Bio-Rad) was performed to determine the number of PCR cycles corresponding to the amplification plateau. Each qPCR included 1 ng of assembled library (S2), 2.5 µL of 10 µM of biotinylated forward primer (skpp504F), 2.5 µL of 10 µM of biotinylated reverse primer (skpp504R) (**Table S2**), 0.25 µL of 100X Biotium Thiazole Green (Thermo Fisher Scientific), 12.5 µL of 2X Q5 High-Fidelity Master Mix (NEB), and nuclease-free water to a total volume of 25 µL. The qPCR program included initial denaturation at 95 °C for 3 min, followed by 40 cycles of 98 °C for 15 seconds, annealing at 65 °C for 30 seconds, and extension at 72 °C for 15 seconds.

In total, sufficient PCRs were prepared to yield 5-10 µg of biotinylated library S4 for downstream use. Bulk amplification PCRs were prepared using the same setup as described for preparing the qPCRs. A 1 min extension step at 72 °C was added to the thermocycler PCR protocol used for qPCR and library S4 was amplified for 12 cycles. The amplified products were purified with 5 µg DNA Cleanup Columns (NEB).

The 1, 536-gene DHFR (S2) fragments were PCR-amplified to add standard Illumina adapters and sequenced on a MiSeq. Amplicons were subsequently re-amplified using biotinylated P5 and P7 primers. To determine the optimal number of PCR cycles, qPCR reactions were prepared using 1 ng of library S2 amplicons, 1.25 µL of 10 µM biotinylated forward primer (P5_FWD_Biotin_NV), 1.25 µL of 10 µM biotinylated reverse primer (P7_REV_Biotin_NV) (**Table S2**), 0.25 µL of 100× Biotium Thiazole Green (Thermo Fisher Scientific), 12.5 µL of 2× Q5 High-Fidelity Master Mix (NEB), and nuclease-free water to a final volume of 25 µL. The qPCR program consisted of an initial denaturation at 98 °C for 30 s, followed by 40 cycles of 98 °C for 10 s, 70 °C for 30 s, and 72 °C for 30 s.

Bulk amplification was then performed until 5–10 µg of biotinylated DNA was obtained for downstream large-scale diversity generation. Bulk PCR reactions were prepared identically to the qPCR reactions described for library S4, except that the thermocycler protocol included a 2 min final extension at 72 °C. Library S2 was amplified for 24 cycles, and the resulting products were purified using 5 µg DNA Cleanup Columns (NEB).

### Generation of large-scale diversity of barcoded gene libraries

Barcoded gene libraries were digested, ligated into barcoding plasmids, and transformed into electrocompetent *E. coli* with high efficiency to generate large-scale library diversity. This approach was adapted from the methodologies of Plesa and colleagues (15).

First, the pEVBC barcoding plasmids and gene library inserts, such as *mCherry* or DHFR libraries, were digested with KpnI-HF® and NdeI restriction enzymes to generate compatible sticky ends. Each digest contained 666 ng of pEVBC backbone or 333 ng of gene insert, 5 µL of 10X CutSmart® Buffer, 1 µL of KpnI-HF® (20, 000 units/mL) (NEB), 1 µL of NdeI (20, 000 units/mL) (NEB), and nuclease-free water to a total volume of 25 µL. The pEVBC digests also included 1 µL of Shrimp Alkaline Phosphatase (rSAP, 1, 000 units/mL) (NEB) to prevent plasmid recircularization. Digestions were incubated at 37 °C for 1.5 hours, followed by heat inactivation of NdeI and rSAP at 65 °C for 20 minutes. The digested products were then purified using 5 µg DNA Cleanup Columns (NEB).

Next, uncut biotinylated molecules were removed with streptavidin magnetic beads to concentrate digested products. Streptavidin magnetic beads (NEB, Ipswich, USA), ranging from 30 to 60 µL at a concentration of 4 mg/mL, were prepared by removing the storage buffer and performing equilibration washes according to the manufacturer’s instructions. The quantity of beads used per digest was determined based on their binding capacity of 1 mg per 500 pmol of 25 nt single-stranded DNA, as specified by the manufacturer. This amount was adjusted according to the size and quantity of the backbone or insert. The equilibrated beads were then resuspended with twice the original volume of 2X binding and wash buffer (2X B&W buffer; 2M NaCl, 1mM EDTA, 10mM Tris) and added to the digested DNA in 1.5 mL tubes. The bead capture samples were incubated on a thermomixer at RT for 30 min with shaking at 1700 RPM.

After incubation, a DynaMag-2 magnetic rack was used for 1 min to separate the beads. The supernatant fraction was transferred to a clean 1.5 mL tube, and the beads were subsequently resuspended in 50 µL of 2X B&W buffer. The magnetic bead capture process was repeated three times. The supernatant fractions containing cut non-biotinylated DNA were purified with 5 µg DNA Cleanup Columns (NEB).

The digested pEVBC backbones and gene library insert sticky ends were ligated together using T4 DNA ligase at a 1:4 plasmid to insert ratio. For each gene library, 4-6 ligation reactions were prepared. Each full ligation (FL) assembly reaction consisted of 0.05 pmol of digested pEVBC backbone combined with approximately 0.2 pmol of gene library insert, except for the no insert control (NIC). The ligation mixtures included 4 µL of 10X T4 DNA ligase reaction buffer (NEB) supplemented with a final concentration of 1 mM ATP (New England Biolabs, NEB, Ipswich, USA), 2 µL T4 DNA ligase (400, 000 units/mL) (NEB, Ipswich, USA), and nuclease-free water to a final volume of 20 µL. The ligations were incubated overnight (∼22 hours) at 16 °C in a thermocycler, followed by purification using 5 µg DNA Cleanup Columns (NEB). The ligated products were eluted in the minimum volume of Monarch® DNA Elution Buffer (NEB) possible for the DNA columns, typically 7 µL, to achieve a final concentration between 50 and 100 ng/µL.

To remove guanidinium salt contaminants from FL and NIC products, drop dialysis was performed. Each ligated gene library was pipetted onto a 50 nm pore nitrocellulose membrane floating on deionized water contained in an 150 x 15 mm petri dish (Kord-Valmark Labware Products, Bristol, USA). After 20 minutes of incubation, each ligation was transferred to a clean 1.5 mL tube and quantified.

The ligated products were then transformed into *E. coli* to generate high-diversity libraries. FL and NIC ligated products were initially transformed into NEB® 5-alpha Electrocompetent *E. coli*, and later into NEB® 10-beta Electrocompetent *E. coli* once the 5-alpha cells were discontinued (NEB). Electrocompetent cells were thawed and 25 µL aliquots were distributed to 1.5 mL tubes chilled on ice. One to two microliters of 50-100 ng of ligated products were added to each aliquot of cells. Three cell aliquots received each individual FL product, while NIC and the supercoiled pUC19 (Bayou Biolabs, Metairie, USA) transformation positive control were each added to separate aliquots. After flicking each 1.5 mL tube to mix the DNA and cells, the mixture was loaded onto Gene Pulser/MicroPulser Electroporation Cuvettes (Bio-Rad Laboratories Inc, Hercules, USA). After BioRad Micropulser electroporation, the cells were resuspended in Super Optimal Broth with Catabolite Repression Medium (NEB) and incubated at 37 °C for 1 hour in an Innova shaking incubator.

All three recovered FL transformants, initially diluted to 1:10 from the original cell aliquots, were combined. A 30 µL aliquot of FL transformants was then diluted 10-fold to prepare a 1:100 dilution. This 1:100 dilution underwent four additional 10-fold serial dilutions to prepare colony counting plates. Similarly, a dilution series was prepared for the single aliquot of pUC19 transformants. In contrast, 100 µL of NIC transformants were serially diluted 10-fold and 100-fold to achieve 1:100 and 1:1000 dilutions, respectively. Finally, 100 µL of each prepared dilution was spread-plated in duplicate on Lysogeny broth (LB) agar supplemented with 0.1 mg/mL carbenicillin. Additionally, large-scale transformed libraries were plated by adding 300 µL of 1:10 dilution FL transformants to 5-10 pre-warmed 150 x 15 mm plates containing LB and 0.1 mg/mL carbenicillin. All plates were then incubated overnight at 37 °C.

Colonies from the dilution series were counted to determine CFU/mL for FL, NIC, and pUC19 transformations. The FL dilution series yielded at least 1.0 x 10^6 CFU/mL, with large-scale transformations showing lawn growth, indicating sufficient diversity in the transformed FL gene libraries. Additionally, the NIC CFU/mL, consistently lower by several orders of magnitude compared to FL, suggested minimal background from self-ligated pEVBC backbone.

To harvest FL gene library bacterial lawns, 5-10 mL of LB was dispensed onto each large 150 x 15 mm plate and the bacterial lawns were scraped with a sterile spreader. The scraped lawns were transferred into a 50 mL tube and the OD600 was measured. Multiple glycerol stocks of both undiluted and diluted (OD 4-7) cells were stored at -80 °C. The remaining diluted cells were partitioned into 5 mL aliquots, spun down into pellets, and stored at -80 °C. Supercoiled gene libraries were isolated from the scraped cells by thawing one or two pellets and using the Monarch® Plasmid Miniprep Kit protocol or a midiprep kit.

### Indexed amplification of the barcoded rfp gene library for MiSeq

Primers designed to add Illumina adapters and a sequencing index to the barcoded library were ordered from Integrated DNA Technologies (IDT, Coralville, USA). The forward primer (mi7_FWD_Amp_NV) annealed to the 5’ end of the barcode, incorporating a sequence buffer and P5 Illumina adapter. The reverse primer (mi7_REV_Amp4_NV) annealed 458 bp downstream of the barcode, adding a P7 Illumina adapter, sequence buffer, and Illumina N701 index 1 (**Table S2**). The resulting PCR product was 586 bp and included the barcode without *rfp*.

qPCR (CFX Opus 96, Bio-Rad) was performed to determine the optimal number of cycles needed to amplify the barcode region of the *rfp* library. Each qPCR was prepared with 1 ng of scraped *rfp* library, 2 µL each of 100 nM forward and reverse primers, 10 µL of 2X Kapa Fast SYBR Green (Roche), and nuclease-free water to 20 µL. The qPCR program included initial denaturation at 98 °C for 30 seconds, followed by 40 cycles of 98 °C for 10 seconds, annealing at 60 °C for 30 seconds, and extension at 72 °C for 20 seconds.

Ten PCRs were prepared to amplify the barcode product for sequencing. Each PCR contained 1 ng of scraped *rfp* library, 2.5 µL of 10 µM forward and reverse primers, 25 µL of 2X Q5 High-Fidelity Master Mix (NEB), and nuclease-free water to 50 µL. The thermocycler protocol included a 2-minute extension at 72 °C, with amplification proceeding for 24 cycles as determined by qPCR. The amplified products were purified with 5 µg DNA Cleanup Columns (NEB) and size-selected at the 600 bp band from a 2.0% agarose gel using the Monarch® DNA Gel Extraction Kit (NEB). Purified products were submitted to the UO Genomics & Cell Characterization Core Facility (GC3F) for sequencing. Additionally, 30 µL of 100 µM custom sequencing primers for read 1 (Mi7_R1_NV), read 2 (Mi7_R2_NV), and the index read (Mi7_Rindex) were submitted. For Miseq sequencing, 25 million paired-end reads with a 75 bp read length were requested.

### Indexed amplification of the barcoded DHFR gene libraries for MiSeq

We ordered primers from Integrated DNA Technologies (IDT; Coralville, USA) to append Illumina adapters and sequencing indices to two barcoded *DHFR* libraries assembled by DropSynth. The forward primer (mi9_FWD_Amp_NV) annealed to a region adjacent to the 5′ ends of DHFR genes, while reverse primers (mi9_REV_Amp_Index2_NV and mi9_REV_Amp_Index5_NV) incorporated N702 and N705 Illumina indices into the S2 and S4 libraries, respectively (**Table S2**). The resulting PCR products were between 600 and 700 bp in length since they included DHFR genes of various lengths and barcodes.

qPCR (CFX Opus 96, Bio-Rad) was performed on all four libraries to optimize cycle numbers for DHFR gene and barcode amplification. Each qPCR was prepared with 1 ng of scraped pEVBC8-DHFR library,

2.5 µL each of 10 µM forward and reverse primers, 25 µL of 2X Q5 High-Fidelity Master Mix (NEB), and nuclease-free water to 50 µL. The qPCR program included initial denaturation at 98 °C for 30 seconds, followed by 40 cycles of 98 °C for 10 seconds, annealing at 72 °C for 30 seconds, and extension at 72 °C for 60 seconds.

Four PCRs were prepared to bulk amplify each library for sequencing using the same protocol as the qPCR. However, library S4 was amplified for 16 cycles and library S2 for 20 cycles. Amplified products were purified using 5 µg DNA Cleanup Columns (NEB) and size-selected from a 2.0% agarose gel with the Monarch® DNA Gel Extraction Kit (NEB). Purified products and 30 µL of 100 µM custom sequencing primers (Mi9_R1_NV, Mi9_R2_NV, Mi9_EVBC8_Rindex) (**Table S2**) were submitted to Admera Health for 2 x 300 bp paired-end sequencing reads. Due to low Miseq sequencing depth, samples were resubmitted for sequencing.

### Illumina library sequencing

We mapped barcodes to gene variants using Illumina Miseq runs analyzed with a custom pipeline described elsewhere (16). Briefly, reads were merged using bbmerge (bbmap 38.18), and a custom python script was used to extract barcodes and variable gene regions. Barcodes were collapsed using Starcode spheres algorithm on distance of 1 (87). A majority call was used to determine the consensus sequence linked to each barcode when it had multiple reads. All required scripts are available on the lab’s GitHub repository (https://github.com/PlesaLab/BC_Mapping).

### sgRNA spacer selection pipeline

The computational spacer selection pipeline converts long-read amplicon sequencing data from a barcoded gene library into a set of single-guide RNAs (sgRNAs) for CRISPR-dCas9 that (i) cover every targeted gene, (ii) avoid recognizing any non-target sequence and (iii) are normalized so that each gene targeted by an sgRNA has a similar number of sequencing reads (**Fig. S9**). The code is written in R 4.3.0 with several small helper scripts in Python. Sequencing reads are first mapped to their corresponding 20-nucleotide barcodes as described previously and collapsed with Starcode (spheres distance ≤ 1) to remove PCR and sequencing errors (87). Each mapped insert is then expanded by appending 20bp of the constant sequences that flank the variable region in the corresponding vector, recreating the entire cloning context. This allows detection of spacers spanning the junction between the constant and variable regions. The full plasmid sequence itself is also loaded so that spacers targeting backbone DNA can be excluded at a later step. Barcodes whose corresponding translated open reading frame exactly matches an entry in the gene design file are provisionally labelled good; all others, together with barcodes identified in the vector or those with undetermined N bases, are labelled bad. We also keep track of the abundance of each unique barcode in the pooled library. For every read, the code searches for all occurrences of the sequence motif *N_20_NGG* where *NGG* represents the canonical SpCas9 PAM. A sliding-window regular expression returns every overlapping 20-nucleotide protospacer immediately upstream of an *NGG*, on both strands. The corresponding barcode sequence is appended to the protospacer list and labeled as either inside the gene region or directly on the barcode. Off-target potential is judged by a 16-nucleotide (adjustable) seed defined as the 3′ end of each 20-mer spacer. We track two kinds of collisions. A seed collision occurs when the same 16-mer is found in at least one good and one bad read. A full-length collision is recorded when an entire 20-mer is shared between the two classes. Protospacers involved in full-length collisions are immediately discarded; seed collisions can be retained or removed in later filtering steps depending on the desired stringency. Because sgRNAs are ultimately cloned by Golden Gate assembly, any protospacer with flanking context sequence which recreates a BsaI recognition site is excluded. A barcode is designated good if it possesses at least one protospacer that is free of full-length collisions, passes the BsaI filter, and if required, is free of 16-mer seed collisions.

Barcode counts within a gene can vary by more than two orders of magnitude and could potentially bias dCas9 capture. To target molecules with similar abundance, the pipeline selects up to three barcodes per gene whose abundance is closest to a single global target value. First, the median barcode read count is calculated for every gene. The medians from all genes are then combined and the median of that distribution is taken as the desired global target count. Within each gene the absolute difference between every barcode’s read count and this target is computed. Barcodes whose counts exactly match the target are accepted directly. If fewer than three such barcodes exist, the remaining positions are filled by barcodes in ascending order of their absolute deviation from the target until either three barcodes have been chosen or no candidates remain. The effectiveness of this normalization is evaluated by plotting Lorenz curves and computing the Gini Coefficient. In the test 1, 384 DHFR dataset, the in-silico Gini Coefficient dropped from 0.63 before normalization to 0.14 afterwards. The complete codebase, including the small Python utilities, is available at the Plesa Lab GitHub repository (https://github.com/PlesaLab/).

### Preliminary exploration of buffer compositions for magnetic bead washes following BAR-CAT enrichment

Before establishing the BAR-CAT v0.1 protocol, we conducted preliminary studies to assess how the number of bead washes and the composition of wash buffers affected stringency following BAR-CAT enrichment. Various buffer conditions were tested using streptavidin magnetic beads. Full experimental details are available in the **Supplementary Methods**.

### BAR-CAT v0.1: Proof-of-concept enrichment of 18 targeted barcodes from the rfp library

CRISPR ribonucleoproteins (RNPs) were assembled using an 18-plex sgRNA library *in vitro* transcribed elsewhere (46). Each RNP assembly reaction contained 3.0 µL of 10X NEBuffer r3.1 (New England Biolabs), 3.0 µL of 300 nM sgRNAs (final: 33.3 nM), 1.0 µL of 1 µM dCas9-3xFLAG-Biotin (Sigma-Aldrich; final: 33.3 nM), 0.75 µL of Murine RNase inhibitor (40, 000 U/mL, NEB), and nuclease-free water to a final volume of 27 µL. Reagents were added in the order listed. Reactions were mixed, pulse-spun, and incubated at 25 °C for 10 min, followed by 10 min at 37 °C in a thermocycler. RNPs were used immediately for downstream enrichment.

To each RNP formation reaction, 1.0 µL (50 ng) of supercoiled pEVBC-ligated gene library was added, and the total volume was adjusted to 30 µL with nuclease-free water. A negative control was prepared by adding 3.0 µL of water to an RNP-only reaction. Reactions were pulse-spun and incubated at 37 °C for 15 min.

Streptavidin magnetic beads (NEB, 4 mg/mL) were equilibrated per manufacturer’s protocol and resuspended in 2X B&W buffer (10 mM Tris-HCl pH 7.5, 1 mM EDTA, 2 M NaCl). To each enrichment reaction, 10 µL of equilibrated beads were added. Capture of dCas9 was performed at 37 °C for 30 min in a thermomixer at 1700 rpm. Beads were collected on a DynaMag-2 rack (Thermo Fisher Scientific), and the supernatant was discarded. Beads were washed six times with 50 µL of 2X B&W buffer, transferring the beads to fresh tubes before the sixth wash. Beads were resuspended in 60 µL of nuclease-free water and stored at 4 °C prior to PCR.

To estimate optimal amplification cycles, duplicate 50 µL qPCRs were prepared for each condition. Reactions contained 10 µL of washed beads, 2.5 µL of 10 µM forward (Mi7_FWD_Amp_NV) and reverse (Mi8_REV_Amp4_Index2_NV) primers (**Table S2**), 0.5 µL of 100X Biotium Thiazole Green (Thermo Fisher), 25 µL of 2X Q5 High-Fidelity Master Mix (NEB), and water to 50 µL. Thermocycling was performed on a CFX Opus 96 machine (Bio-Rad) using the following program: 98 °C for 30 s; 40 cycles of 98 °C for 10 s, 72 °C for 30 s, and 72 °C for 2 min.

Bulk amplification of enriched DNA was carried out using the same conditions and number of cycles determined by qPCR. Amplified products were cleaned using Monarch® 5 µg DNA Cleanup Columns (NEB). The 586 bp enrichment products were size-selected using 0.8% SYBR Safe E-Gels (CloneWell™ II, Thermo Fisher Scientific) with a 1 kbp Plus DNA Ladder (NEB) as reference. Final gel-purified DNA was concentrated using new Monarch® 5 µg Cleanup Columns and submitted to Plasmidsaurus (Eugene, USA) for nanopore sequencing.

### BAR-CAT v0.2: Enrichment of 18 targeted barcodes from the rfp library with optimized bead washes

To improve removal of non-enriched barcodes, BAR-CAT v0.2 incorporated increased wash volumes and additional wash steps following dCas9 capture by streptavidin magnetic beads. All reactions used the barcoded *rfp* gene library and were assembled as described for BAR-CAT v0.1, including RNP formation, target DNA addition, and bead capture.

Following bead capture, each reaction was transferred to a 5 mL tube and placed on a 15 mL tube magnetic rack (Sergi Lab Supplies, Seattle, WA, USA) for high-stringency washing. After 1 minute on the rack, the supernatant was removed, and beads were resuspended in 2 mL of 2X B&W buffer (10 mM Tris-HCl pH 7.5, 1 mM EDTA, 2 M NaCl). Beads were incubated for 1 minute per wash. A total of nine washes were performed per sample, with the beads transferred to fresh tubes prior to the sixth wash to reduce carryover. After the final wash, beads were resuspended in 60 µL of nuclease-free water, vortexed at 2700 rpm, and centrifuged briefly to collect the beads. Samples were stored at 4 °C prior to amplification. qPCR, bulk PCR, cleanup, gel extraction, and nanopore sequencing were performed as described for BAR-CAT v0.1.

### BAR-CAT v0.3: Denaturation and removal of dCas9 prior to amplifying enriched DNA

BAR-CAT v0.3 introduced a dCas9 denaturation step after bead washes to release enriched DNA into the supernatant for downstream amplification. This eliminated the need to amplify directly from the beads, reducing the risk of non-specific DNA carryover. We also evaluated whether DNA format affected enrichment by comparing results from linearized and supercoiled versions of the *rfp* barcode library. All enrichments targeted 18 barcodes from the barcoded *rfp* gene library. dCas9 capture by magnetic beads was performed as described for BAR-CAT v0.1.

To generate linearized *rfp* gene library DNA, we first determined the optimal amplification cycle number by qPCR using 10 µM forward (Mi9_FWD_Amp_NV) and reverse (Mi9_REV_amp_NV) primers, as outlined in the section ***Indexed amplification of the barcoded DHFR gene libraries for MiSeq***. Bulk amplification was then performed using 14 cycles to avoid overamplification. The 827 bp PCR products were purified using Monarch® 5 µg DNA Cleanup Columns (NEB) and verified on 1% agarose gels with a 100 bp Plus DNA Ladder (NEB).

For CRISPR enrichment using either linearized or supercoiled *rfp* libraries, bead washes were performed as described for BAR-CAT v0.2: nine washes in 2 mL of 2X B&W buffer using 5 mL tubes on a 15 mL magnetic rack. After the final wash, beads were resuspended in 50 µL of nuclease-free water.

To denature dCas9 and release enriched DNA from supercoiled library samples, we incorporated a proteinase K digestion step. For each reaction, 25–50 µL of dCas9 capture beads were combined with 1 µL of proteinase K (800 U/mL, NEB) in 1.5 mL tubes. Nuclease-free water was added to a final volume of 50 µL. The mixtures were vortexed, spun down, and incubated at room temperature for 10 minutes in a thermomixer. Samples were then placed on a DynaMag-2 magnetic rack (Thermo Fisher Scientific, Waltham, USA), and the supernatant was collected for downstream purification using Monarch® 5 µg DNA Cleanup Columns (NEB). Enriched DNA was eluted in 50 µL of DNA elution buffer and amplified by qPCR and PCR.

Alternative dCas9 denaturation methods were also tested but not adopted in the final BAR-CAT v0.3 protocol due to lower performance. These included: (1) boiling beads at 65 °C for 5 minutes in a thermomixer at 1700 rpm, and (2) treatment with 200 µL of 8 M urea for 30 minutes at room temperature, as described by Aalipour and colleagues (43). For each, the supernatant was collected post-treatment. Urea-treated samples were further purified using Monarch® DNA Cleanup Columns before amplification.

To determine appropriate PCR cycle numbers for enriched DNA, qPCR was performed for all conditions, including negative controls, using Mi9_FWD_Amp_NV and Mi9_REV_amp_NV primers as described in the section ***Indexed amplification of the barcoded DHFR gene libraries for MiSeq***. Bulk amplification was then conducted using cycle numbers derived from qPCR. Final PCR products were purified using Monarch® 5 µg DNA Cleanup Columns (NEB).

Enriched 827 bp products were size-selected using CloneWell™ II Agarose Gels with SYBR Safe 0.8% E-Gels™ (Thermo Fisher Scientific) and a 1 kbp ladder (NEB). In later experiments, 2% agarose gels and the Monarch® Plasmid Miniprep Kit were used for size selection. All enriched products were submitted to Plasmidsaurus (South San Francisco, USA) for nanopore sequencing.

### BAR-CAT v1.0: Optimized enrichment of three target barcodes using increased rfp gene library input

BAR-CAT v1.0 achieved the best barcode enrichment by incorporating a 10-fold increase in input DNA from the barcoded *rfp* gene library, thereby providing more target molecules for RNP binding. Enrichments were performed using both the supercoiled *rfp* gene library and later barcoded DropSynth DHFR libraries.

CRISPR RNPs were prepared as previously described, using three pooled synthetic sgRNAs (Integrated DNA Technologies) for enrichment from the *rfp* gene library, or a single synthetic sgRNA or *in vitro* transcribed sgRNA libraries prepared as described by Villegas and colleagues (46) for DropSynth DHFR enrichment.

To perform BAR-CAT enrichment, 500 ng of supercoiled barcoded gene library (up to 3 µL) was added to 27 µL of RNPs in qPCR tubes, then brought to a final volume of 30 µL with nuclease-free water. A negative control reaction contained 3 µL of water in place of library DNA. All reactions were mixed, pulse-spun, and incubated at 37 °C for 15 minutes.

For dCas9 pull-down, 5 µL of streptavidin magnetic beads (per reaction) were equilibrated with 2X B&W buffer at twice the final volume. Reactions were transferred from qPCR tubes to 1.5 mL microcentrifuge tubes, and 10 µL of equilibrated beads were added to each. These dCas9-capture reactions were incubated in a thermomixer at 37 °C with shaking at 1700 rpm for 30 minutes.

Bead-bound reactions were transferred to 5 mL tubes for high-stringency washing. Each tube was placed on a 15 mL magnetic rack (Sergi Lab Supplies, Seattle, USA) for one minute to pellet the beads.

Supernatant was discarded, and beads were resuspended in 2 mL of 2X B&W buffer for a one-minute incubation. Each reaction underwent nine such washes; before the sixth wash, beads were transferred to fresh 5 mL tubes. After the final wash, beads were resuspended in 50 µL of nuclease-free water, vortexed at 2700 rpm, spun briefly, and stored at 4 °C until elution.

For proteinase K digestion, 25–50 µL of dCas9-bound beads per sample were combined with 1 µL of Proteinase K (800 U/mL, NEB) and nuclease-free water to a total volume of 50 µL. Tubes were mixed, spun down, and incubated for 10 minutes at room temperature in a thermomixer. Afterward, samples were placed on a DynaMag-2 magnetic rack (Thermo Fisher Scientific, Waltham, USA), and the supernatant containing the enriched DNA was collected and purified using either Monarch® (NEB) or GeneJET (Thermo Fisher Scientific) 5 µg DNA Cleanup Columns. DNA was eluted in 20 µL of elution buffer.

To determine the optimal number of amplification cycles, qPCR was performed on all enrichment samples, including the negative control, using 10 µM Mi9_FWD_Amp_NV and Mi9_REV_amp_NV primers and 2 µL of enriched DNA as template. Reactions followed the protocol for ***Indexed amplification of the barcoded DHFR gene libraries for MiSeq***, with a shortened extension time of 30 seconds.

Bulk PCRs were then prepared for each condition using the same setup and cycle numbers identified by qPCR. Amplified products were purified using Monarch® 5 µg DNA Cleanup Columns and validated for correct size using 0.8%, 2%, or 4% E-Gels™ (Thermo Fisher Scientific). Final enrichment products from both the *rfp* gene library and DropSynth DHFR libraries were submitted to Plasmidsaurus (South San Francisco, USA) for nanopore sequencing.

### Nanopore data analysis and enrichment scores

Nanopore sequencing was used to analyse the enrichment data. Enrichments up to and including testing the different methods to denature dCas9 were run on an R9.4.1 flowcell with v10 chemistry called using Guppy on super-accurate mode. Subsequent enrichments were sequenced with an R10.4.1 flowcell, v14 chemistry and Guppy 6.5.7 on super-accurate mode. Raw fastq data was loaded into a custom python script which extracted the barcode region from each read. Experiments with the pEVBC3 plasmid used the motif ACCTAAGTGTCGCTGCCGAACAGG N_20_ GCTAGAAGAGCGCACGACGTCACG while experiments with pEVBC8 used GGTACCTAAGTGTCGCTGCCGAACAGC N_20_ AGGAGAAGAGCGCACGACGTCACG, where N_20_ is the barcode region. Extracted barcodes were collapsed using the Starcode spheres algorithm on distance of 1 (87). Barcode reads were imported into R for further analysis. We determined the fraction of the population for each barcode by normalizing reads by total sequencing depth. Log_2_ fold enrichment scores were calculated using the ratio of the population fraction after enrichment to their fraction in the initial population. For barcodes not in the initial population a pseudocount of 0.5 was used. Barcodes which dropped out of the population were tracked separately. The population level fold enrichment metric was determined by taking the total population fraction of all targets after enrichment divided by the total fraction of turrets before enrichment. RNA-seq data was analysed as described elsewhere (46). Significance testing was done using a Wilcoxon paired test. CRISPRscan scores were generated using crisprScore (1.4.0) (74).

