## Supplementary Information for "BAR-CAT: Targeted Recovery of Synthetic Genes via Barcode-Directed CRISPR-dCas9 Enrichment"

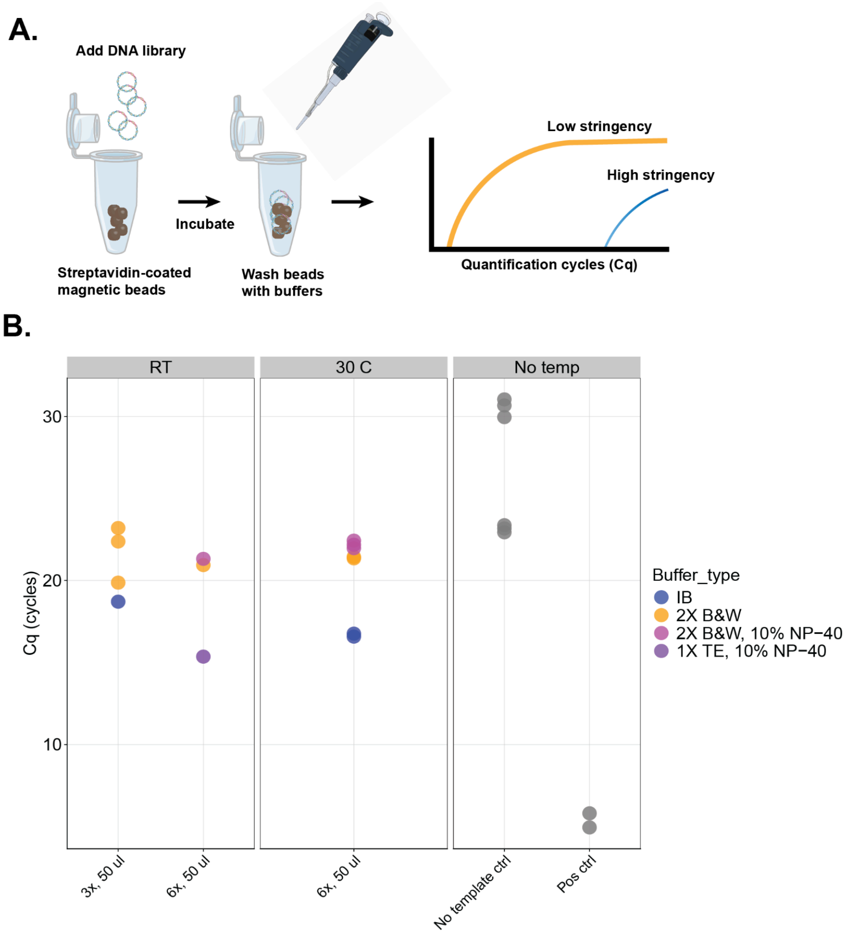

**Figure S1. Wash stringency of various buffers at different temperatures when used to wash streptavidin-coated magnetic beads. A.** Schematic of the bead wash stringency test. Single-gene library DNA was incubated with streptavidin-coated beads in the absence of Cas9 or sgRNAs at 37 °C with shaking. Beads were then washed with one of four buffer formulations (see **Table S1**). Room temperature (RT) washes were performed three times with 50 µL buffer, while washes at 25 °C and 30 °C were performed six times with 50 µL buffer. DNA retention was quantified by qPCR to evaluate the efficiency of nonspecific DNA removal. Icons used: microtube-open-blue by Servier (CC-BY 3.0, https://smart.servier.com/), pipette icon by James-Lloyd (CC0, https://www.badgrammargoodsyntax.com/), and plasmid-l-insert-dna by DBCLS (CC-BY 4.0, https://togotv.dbcls.jp/en/pics.html). **B.** qPCR quantification of DNA remaining on beads following buffer washes under the indicated conditions.

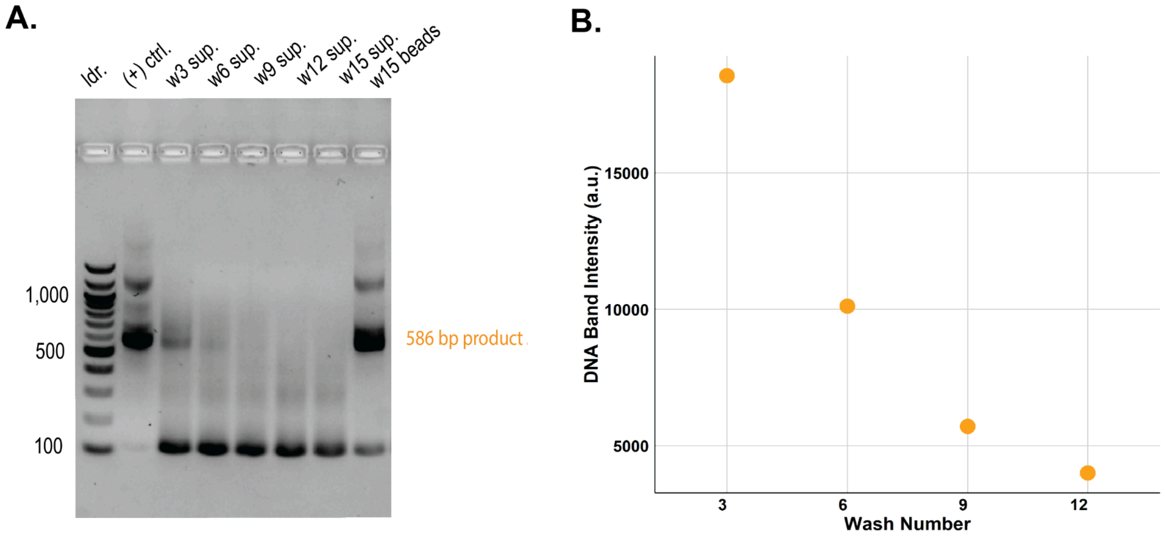

**Figure S2. Evaluating residual DNA after bead washing by performing agarose gel electrophoresis of qPCR products. A.** Agarose gel showing qPCR products. A 100 bp NEB ladder was included for size reference. The positive control contained 448 ng of a 586 bp product amplified from the original supercoiled *rfp* library. SNAP-capture beads were washed 15 times with 1 mL of immobilization buffer (IB, **Table S1**). Supernatants were collected from washes 3, 6, 9, 12, and 15, and the beads were retained after wash 15. qPCR was performed on both the supernatants and the washed beads to amplify any remaining *rfp* library DNA. Bands were visualized using SYBR™ Safe DNA Gel Stain. Presence of the 586 bp product indicates residual DNA not removed by washing. **B.** Quantification of DNA band intensities from the gel in panel **A** using ImageJ (1), shown only for amplification of supernatant samples from washes 3, 6, 9, and 12.

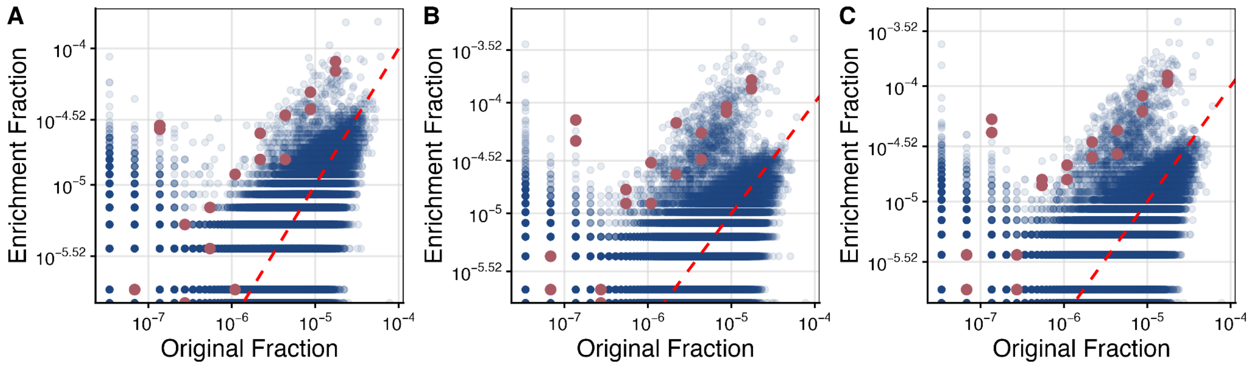

**Figure S3. Barcode distributions before and after enrichment of 18 target barcodes for various bead wash conditions.** Scatter plots comparing the fractional abundance of each barcode in the *rfp* library before (original) and after enrichment under different wash conditions. Each blue dot represents a non-target barcode while each magenta dot represents a barcode targeted by the 18-plex sgRNA library. The red dashed unity line indicates equal representation before and after enrichment and serves as a reference for assessing enrichment. Three bead washing protocols were tested: **A.** Control condition using 6 × 50 µL washes, **B.** 9 × 2 mL washes (used for subsequent enrichments), and **C.** 6 × 5 mL washes.

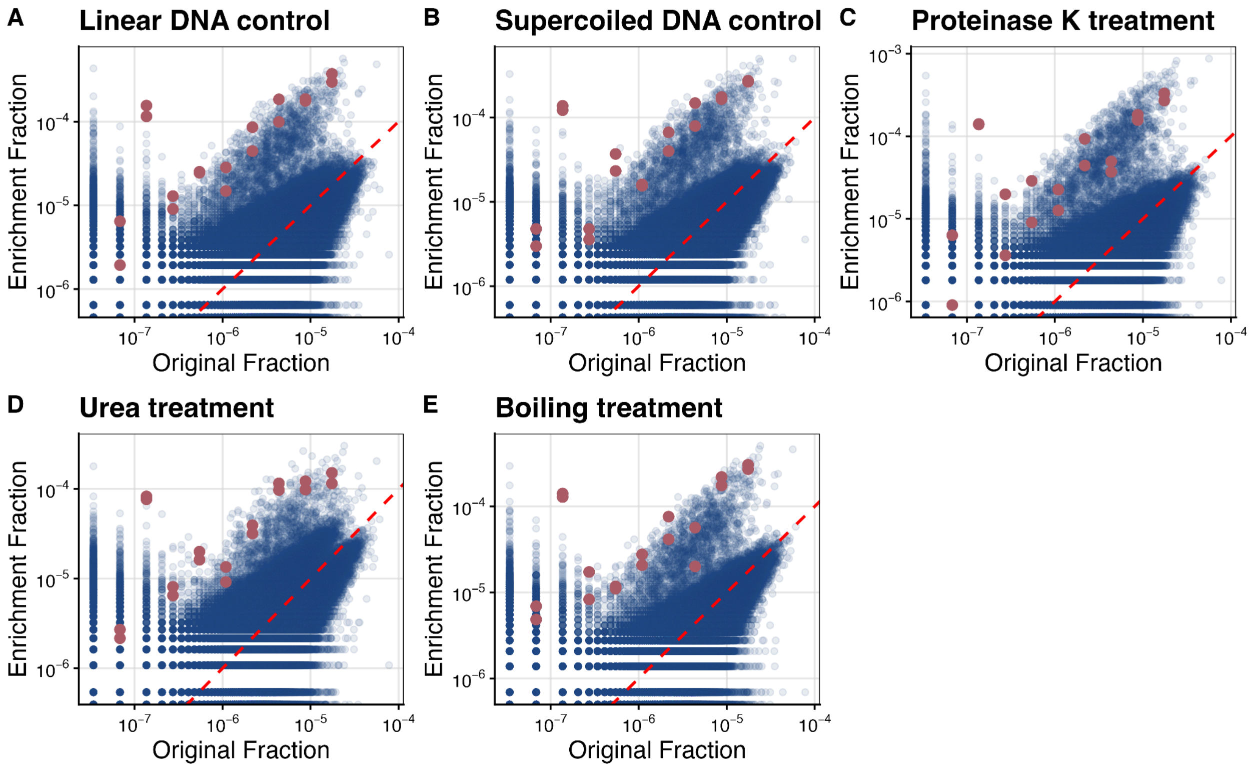

**Figure S4. Barcode distributions before and after enrichment of 18 targeted barcodes for various DNA format and dCas9 denaturation conditions.** Scatter plots comparing the fractional abundance of each barcode in the *rfp* library before (original) and after enrichment under different conditions. Each blue dot represents a non-target barcode, while each magenta dot represents a barcode targeted by the 18-plex sgRNA library. The red dashed unity line indicates equal representation before and after enrichment and serves as a reference for assessing enrichment. The following conditions were tested: **A.** Linear DNA control with amplification off the beads, **B.** Supercoiled DNA control with amplification off the beads, **C.** Supercoiled DNA with proteinase K denaturation of dCas9 and amplification from the supernatant (used for subsequent enrichments), **D.** Supercoiled DNA with 8 M urea denaturation of dCas9 and amplification from the supernatant, **E.** Supercoiled DNA with boiling denaturation of dCas9 and amplification from the supernatant.

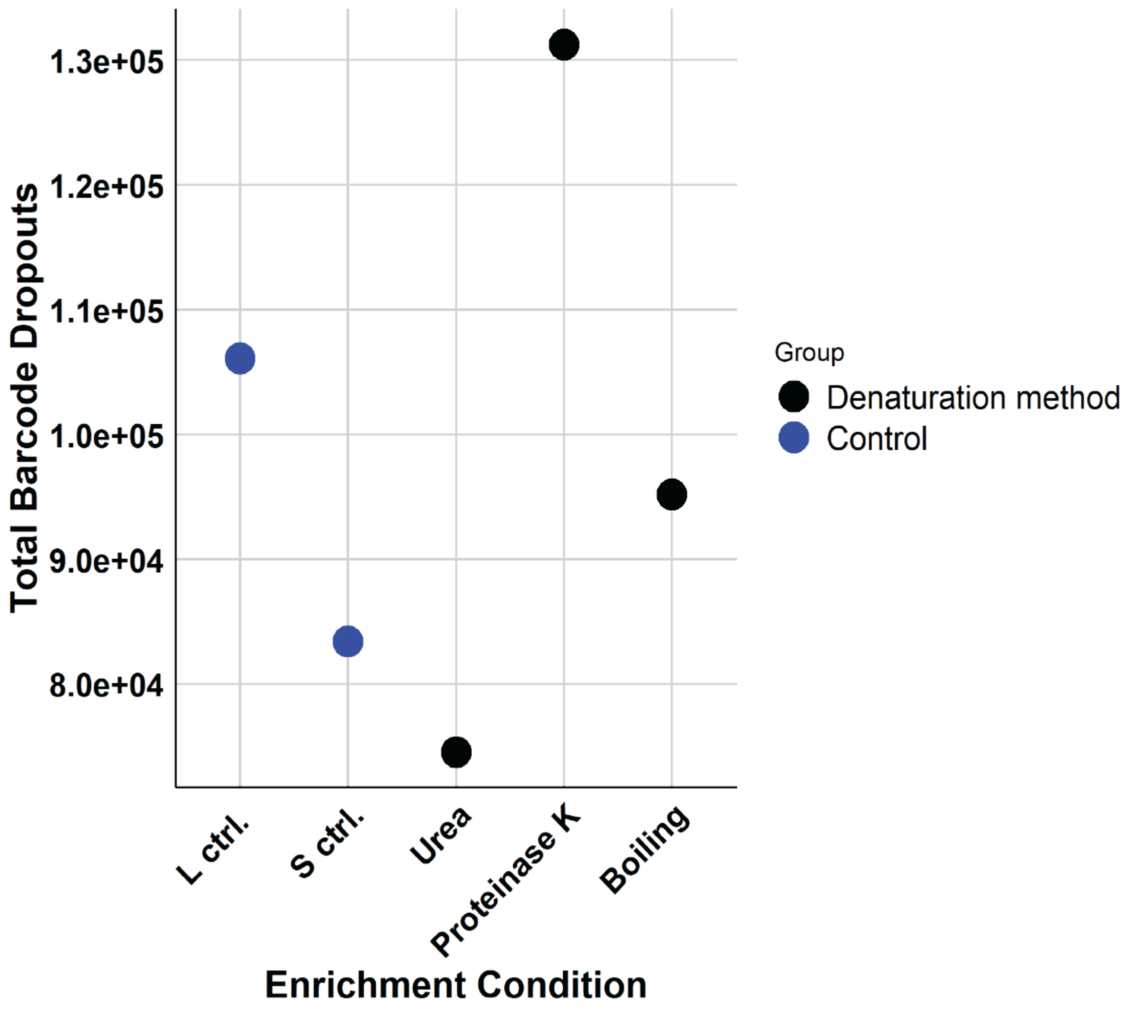

**Figure S5. Total barcode dropout counts are compared across three dCas9 denaturation methods: urea, proteinase K, and heat (boiling).** Two reference controls were included, one using a supercoiled *rfp* library and another using a linearized *rfp* library.

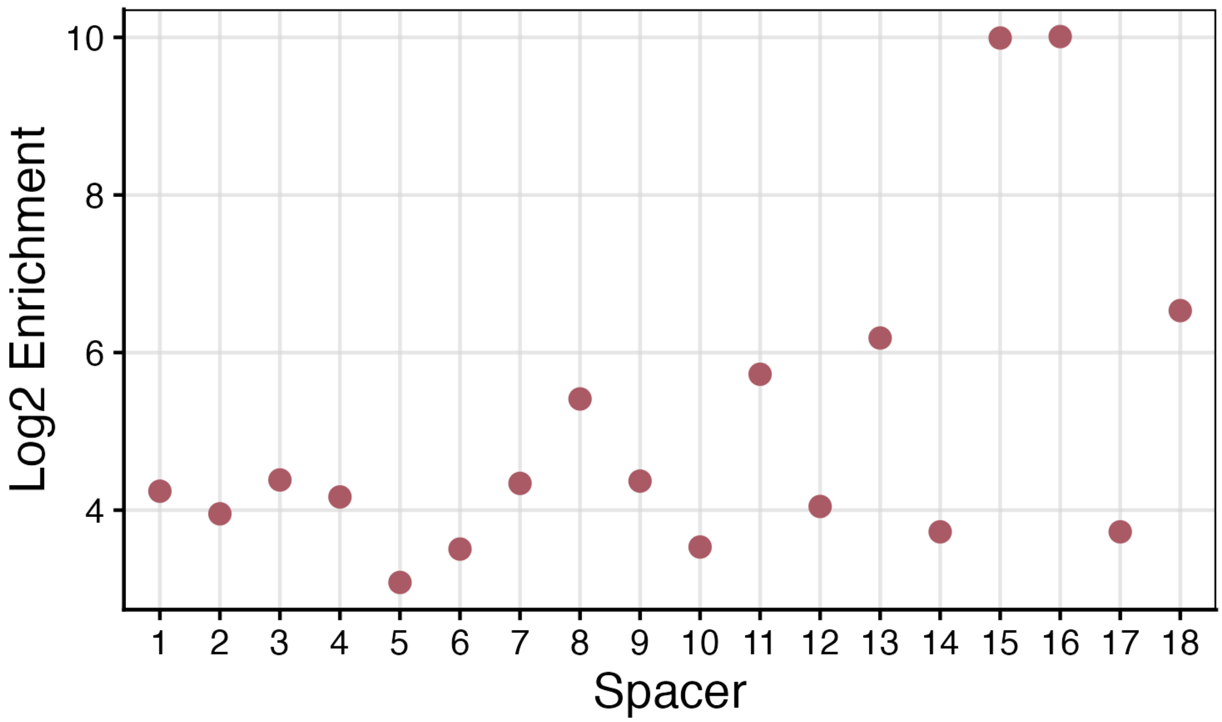

**Figure S6. Log$_2$ enrichment values for the 18 individual barcodes targeted from the *rfp* supercoiled library treated with proteinase K.** The spacers were selected from the 18 barcode protospacers in **Fig. 1B**, with spacer 1 corresponding to the highest abundance barcode and spacer 18 to the least abundant.

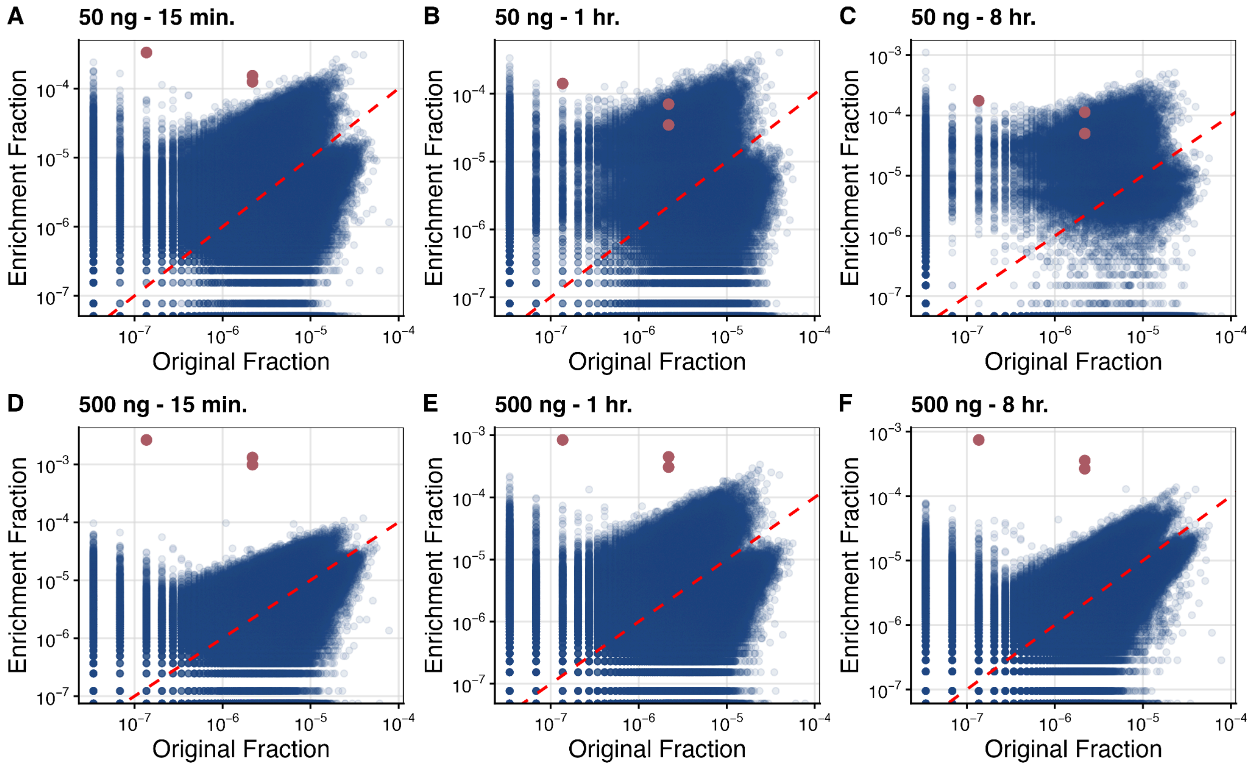

**Figure S7. Barcode distributions before and after enrichment of 18 target barcodes for various DNA input amounts.** Scatter plots comparing the fractional abundance of each barcode in the *rfp* library before (original) and after enrichment under different conditions. Each blue dot represents a non-target barcode, while each magenta dot represents a barcode targeted by three unique synthetic sgRNAs. The red dashed unity line indicates equal representation before and after enrichment and serves as a reference for assessing enrichment. The following conditions were tested: **A.** 50 ng of input DNA with 15 min of enrichment, **B.** 50 ng of input DNA with 1 hour of enrichment, **C.** 50 ng of input DNA with 8 hours of enrichment, **D.** 500 ng of input DNA with 15 min of enrichment, **E.** 500 ng of input DNA with 1 hour of enrichment, **F.** 500 ng of input DNA with 8 hours of enrichment.

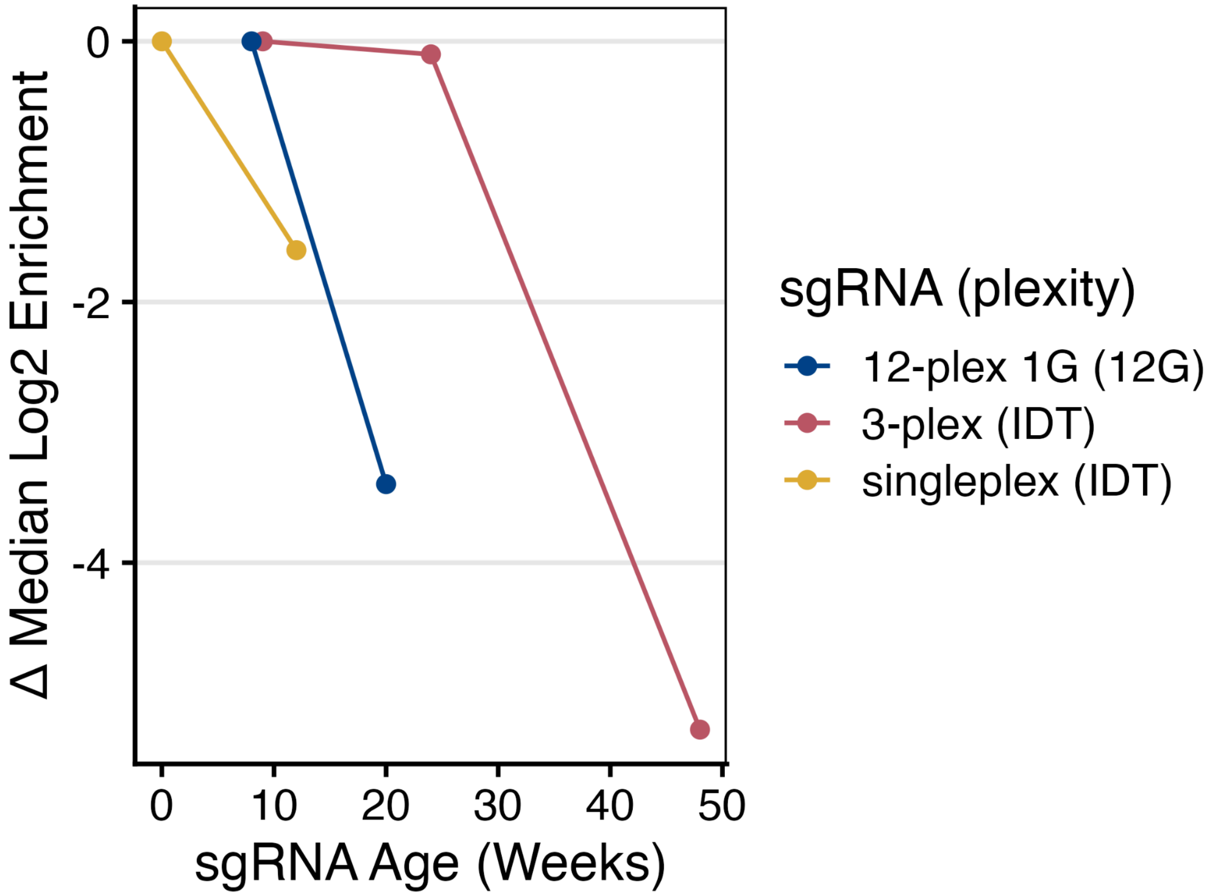

**Figure S8. Change in median $log_2$ enrichment scores as a function of sgRNA age for chemically synthesized and in vitro-transcribed sgRNAs used in BAR-CAT enrichment experiments.** Singleplex and 3-plex pooled sgRNAs (Integrated DNA Technologies) were chemically synthesized, while 12-plex sgRNAs with a 5′ G were generated via *in vitro* transcription using our protocol (2). This analysis evaluates how storage time impacts the performance of sgRNAs used in BAR-CAT.

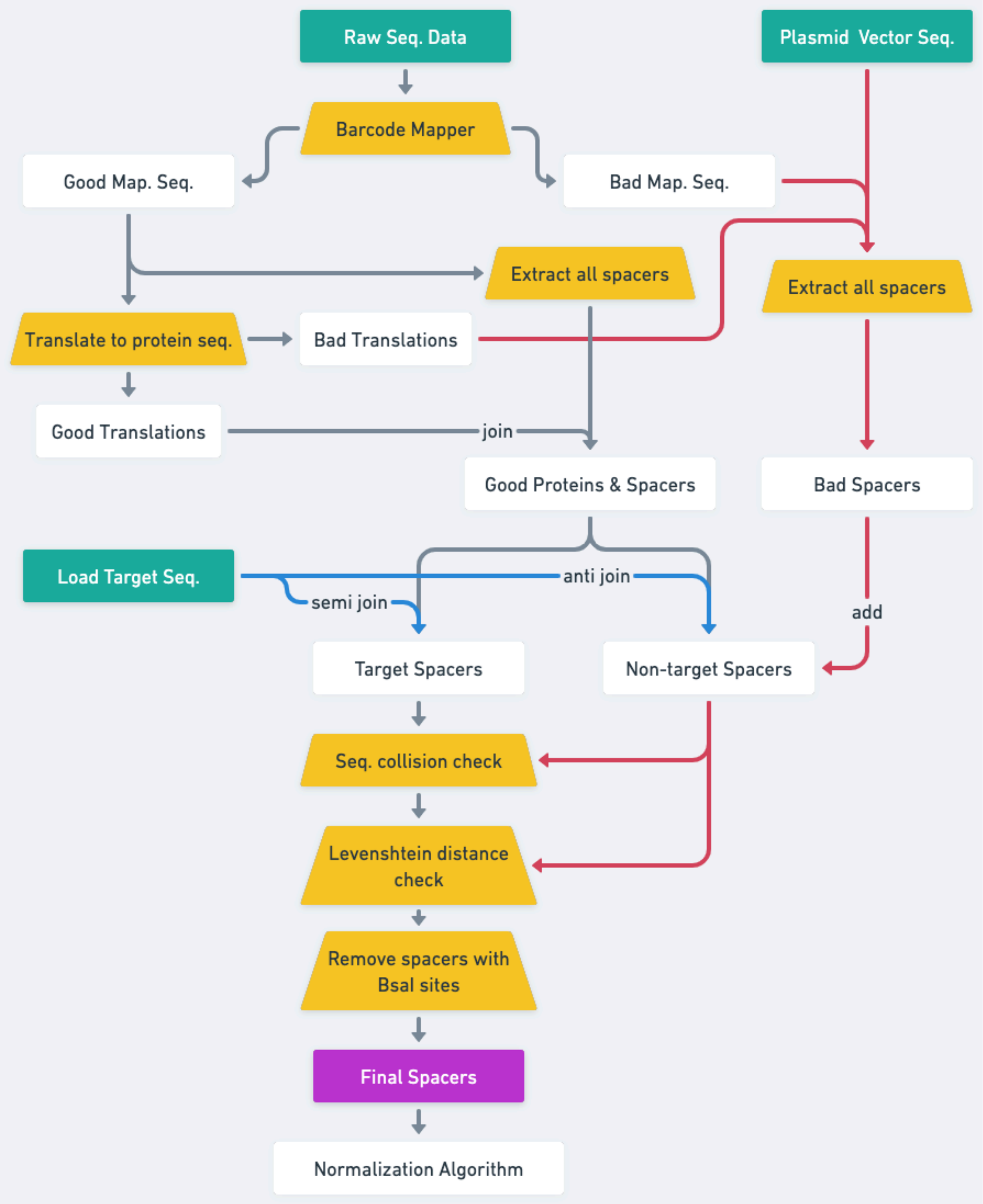

**Figure S9. sgRNA spacer selection pipeline.** Long-read amplicon sequencing data from barcoded gene libraries were transformed into sgRNA libraries using a pipeline. The pipeline was designed to cover every targeted gene, avoid recognizing non-target sequences, and normalized so that the target genes selected had similar numbers of reads. The description of this entire method can be found in the methods section ***sgRNA Spacer Selection Pipeline***.

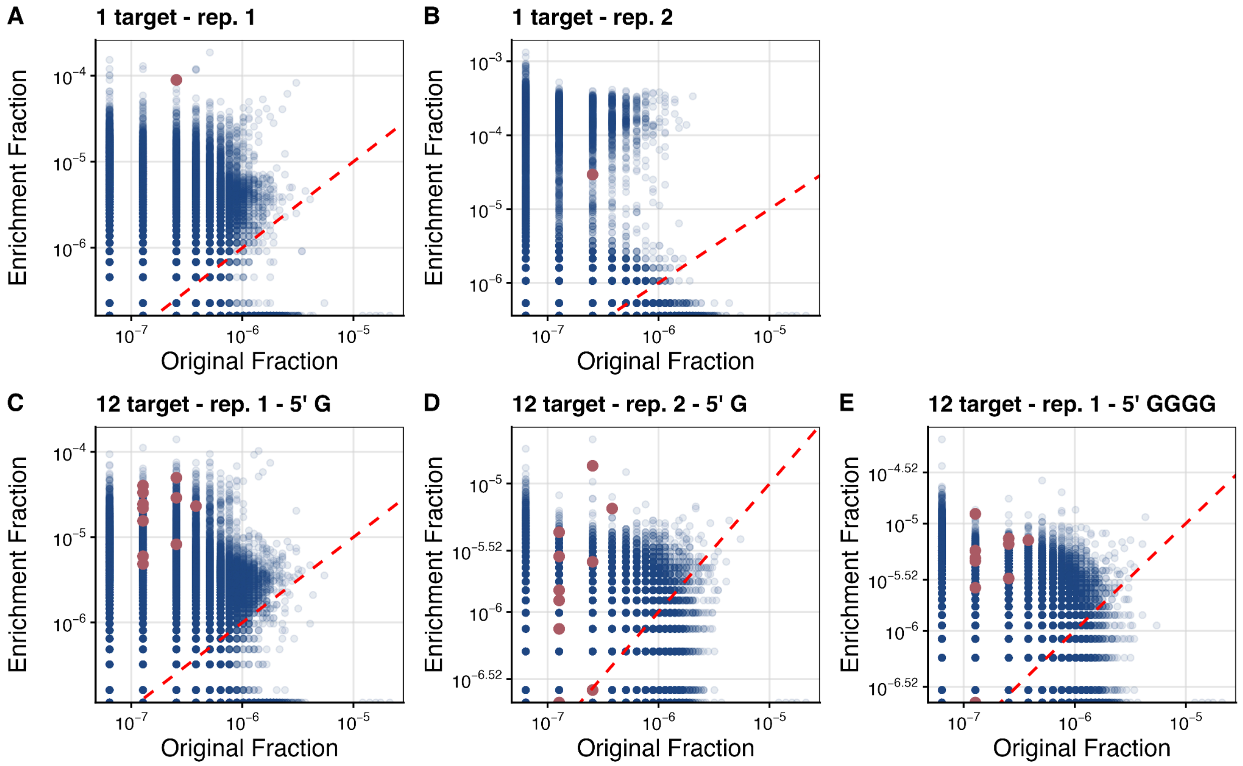

**Figure S10. Barcode distributions before and after singleplex and 12-plex enrichment of a DropSynth dihydrofolate reductase (DHFR) library with varying 5′ guanine additions in sgRNA spacers.** Scatter plots compare the fractional abundance of each barcode in a 384-gene DHFR library (library S4) before (original) and after enrichment. Blue dots represent non-target barcodes; magenta dots indicate target barcodes from either a single synthetic sgRNA with a 5′ guanine (5′ G) or a 12-plex sgRNA pool with either 5′ G or a guanine tetramer (5′ GGGG) at the spacer 5′ end. The red dashed unity line denotes equal representation before and after enrichment, serving as a reference to assess barcode enrichment. Conditions shown: **A.** singleplex enrichment with synthetic 5′ G sgRNA (replicate 1), **B.** replicate 2 of panel A, **C.** 12-plex enrichment with 5′ GGGG sgRNAs (replicate 1), **D.** replicate 2 of panel **C**, **E.** 12-plex enrichment with 5′ GGGG (replicate 1).

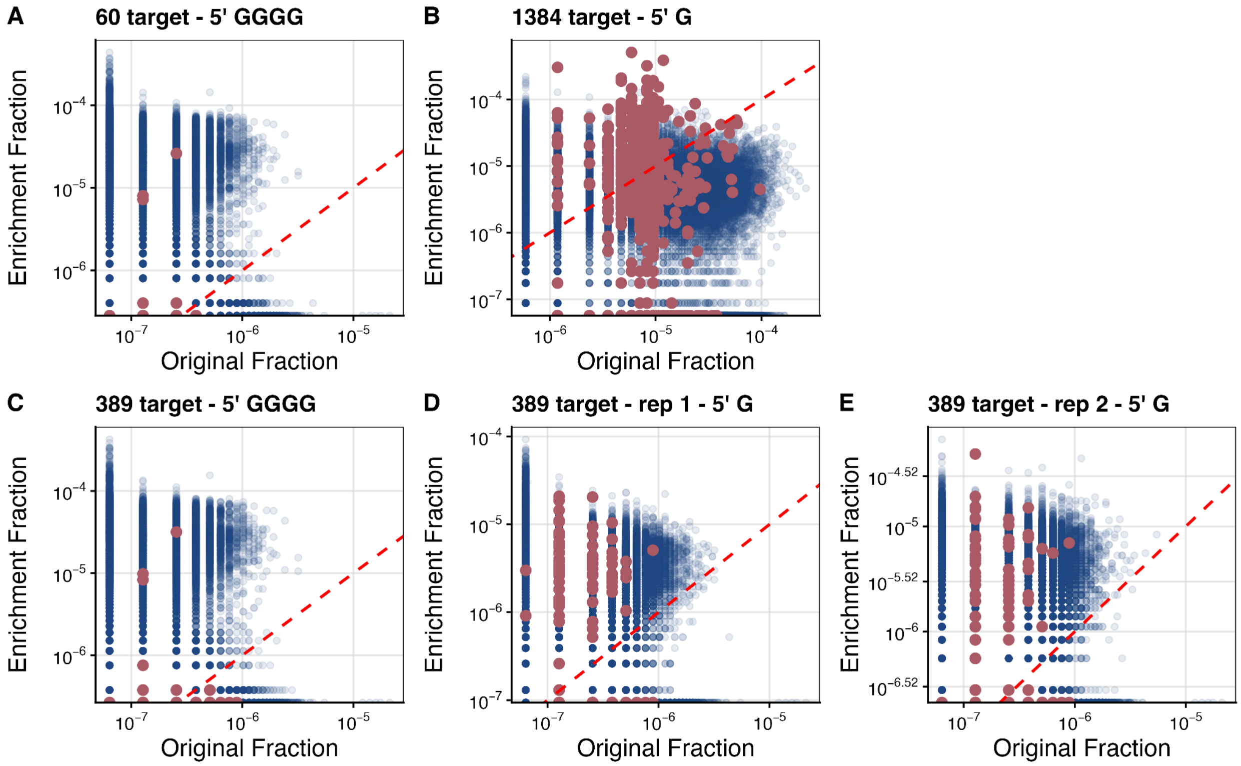

**Figure S11. Barcode distributions before and after 60-, 389-, or 1,384-plex enrichment from DropSynth DHFR libraries with varying 5′ guanine additions in sgRNA spacers.** Scatter plots compare the fractional abundance of each barcode in 384-gene (S4) and 1,536-gene (S2) DropSynth DHFR libraries before (original) and after enrichment. Blue dots indicate non-target barcodes; magenta dots represent target barcodes from in vitro-transcribed sgRNA libraries with spacers beginning with either a 5′ guanine (5′ G) or a guanine tetramer (5′ GGGG). The red dashed unity line marks equal abundance before and after enrichment, serving as a reference to assess enrichment or depletion. Conditions shown: **A.** 60-plex enrichment with 5′ GGGG sgRNAs from a 384 gene DHFR library (S4), B. 1,384-plex enrichment with 5′ G sgRNAs from a 1,536 gene DHFR library (S2), **C.** 389-plex enrichment with 5′ GGGG sgRNAs from a 384-gene DHFR library (S4), **D.** 389-plex enrichment with 5′ G sgRNAs (replicate 1) from a 384-gene DHFR library (S4), **E.** Replicate 2 of panel **C**.

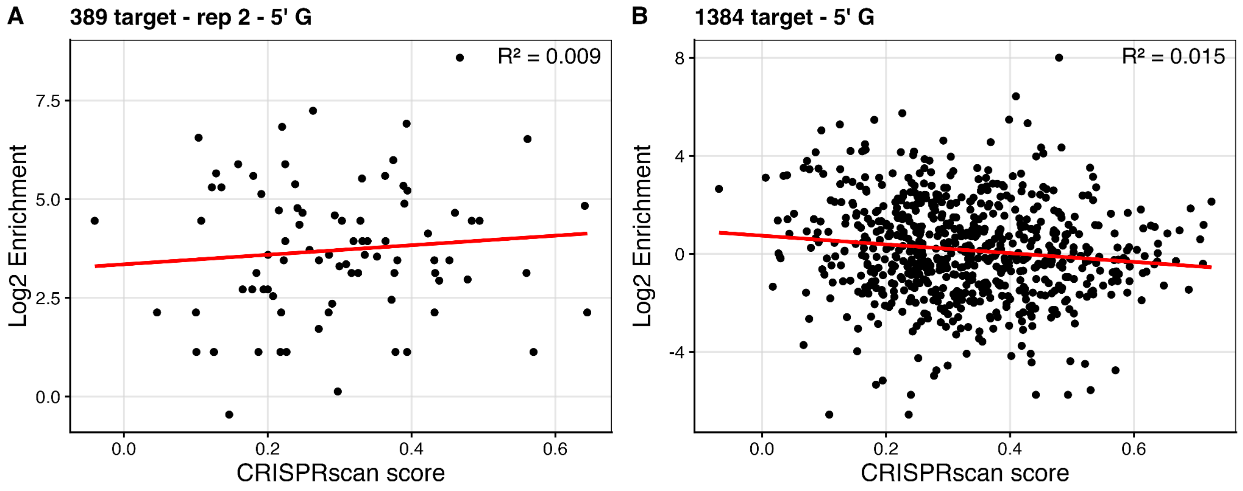

**Figure S12. Correlation analysis of $log_2$ enrichment values for targeted barcodes versus predicted sgRNA performance from the CRISPRscan algorithm** (3)**.** Both the x and y axes are on the log scale and the red line shows the linear regression fit. **A**. Linear regression ($R^2 = 0.009$) for 389-target barcode enrichment with 5′ G sgRNAs (replicate 2) from the 384-gene DHFR library (S4). **B.** Linear regression ($R^2 = 0.015$) for 1,384-target barcode enrichment with 5′ G sgRNAs from the 1,536-gene DHFR library (S2). Other scores are located in this link https://bioconductor.org/packages/release/bioc/vignettes/crisprScore/inst/doc/crisprScore.html

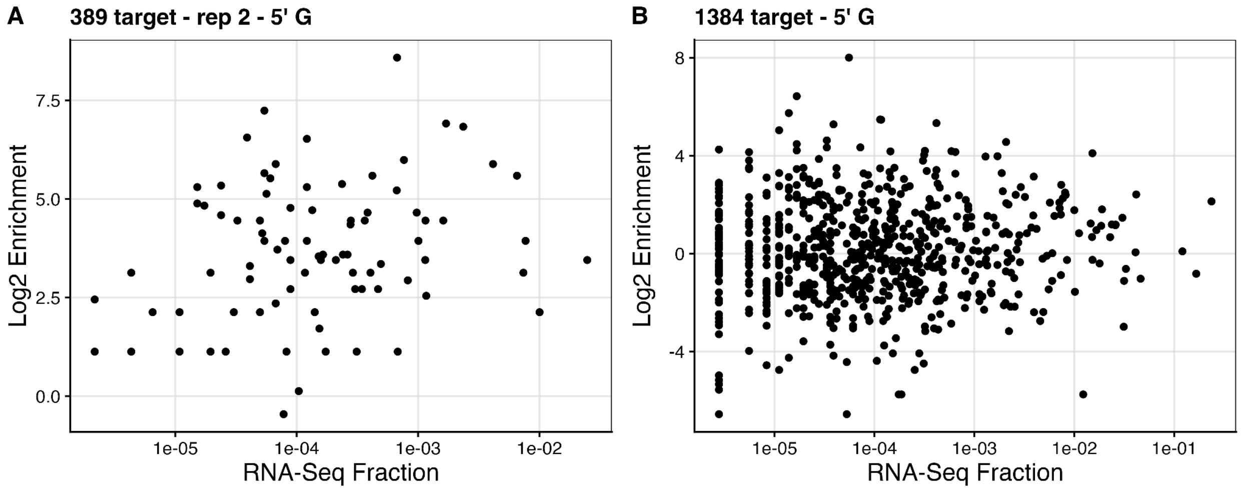

**Figure S13. Distribution of $\log_2$ enrichment values for targeted barcodes relative to the abundance of their corresponding spacers within transcribed sgRNA libraries**. RNA-seq was performed on the sgRNA libraries (2), and the RNA-seq fraction for each spacer was plotted to assess the relationship between enrichment and spacer abundance. **A.** 389-plex enrichment with 5′ G sgRNAs. **B.** 1,384-plex enrichment with 5′ G sgRNAs.

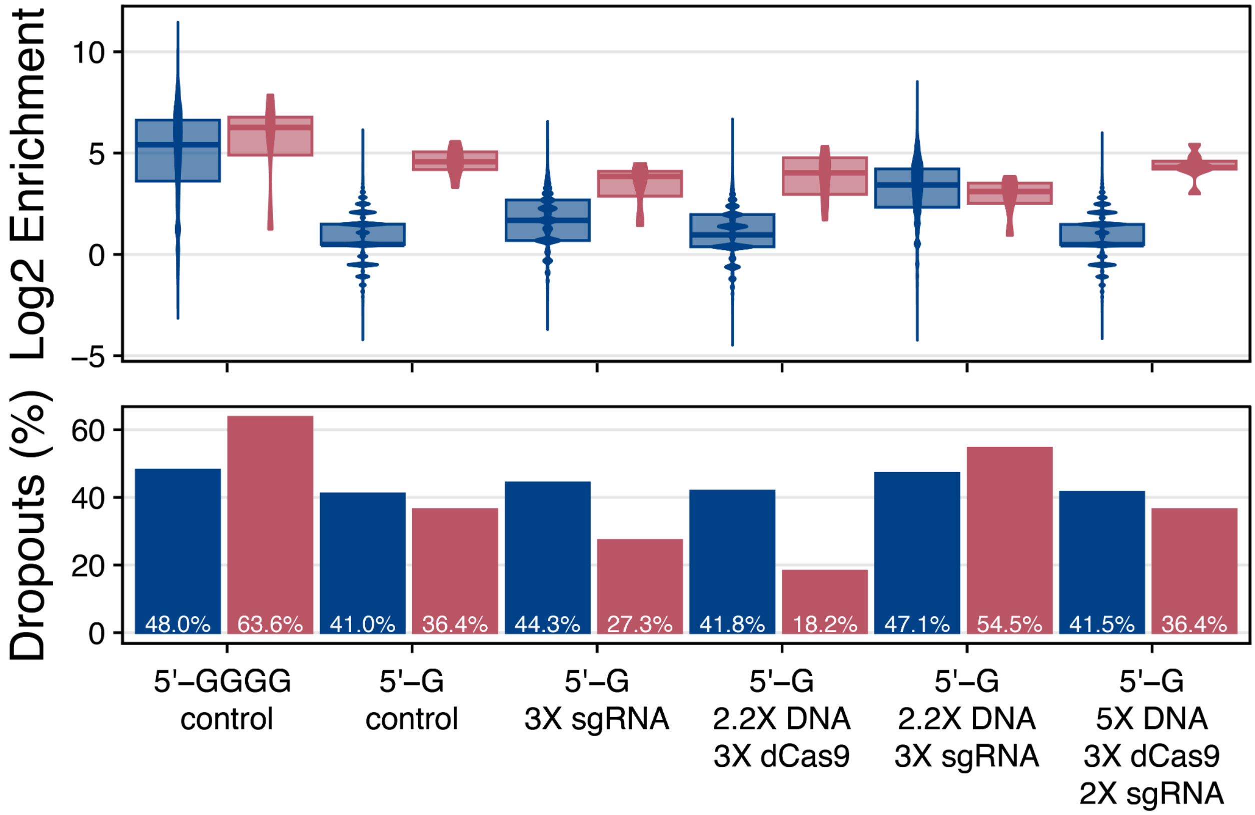

**Figure S14. Assessing the impact of varying DNA, sgRNAs, and dCas9 input amounts on the enrichment of 12 targets from the 384-gene DHFR library (library S4).** Overlaid violin and boxplots showing $\log_2$ enrichment scores, calculated as the $\log_2$ fold change in barcode abundance before and after enrichment. Each distribution compares non-target (blue) and target (magenta) barcodes across the different enrichment conditions tested (x-axis). Shaded areas represent the interquartile range (25th–75th percentile); bars indicate median $\log_2$ enrichment. Percent dropout values are listed in the bar plots corresponding to off-target (blue) and target (magenta) barcodes according to the scale of targeted barcodes, as indicated by the x-axis. None of the modified sgRNA, dCas9, or DNA input amounts affected enrichment scores or barcode dropouts compared to the 5’ GGGG and 5’ G controls.

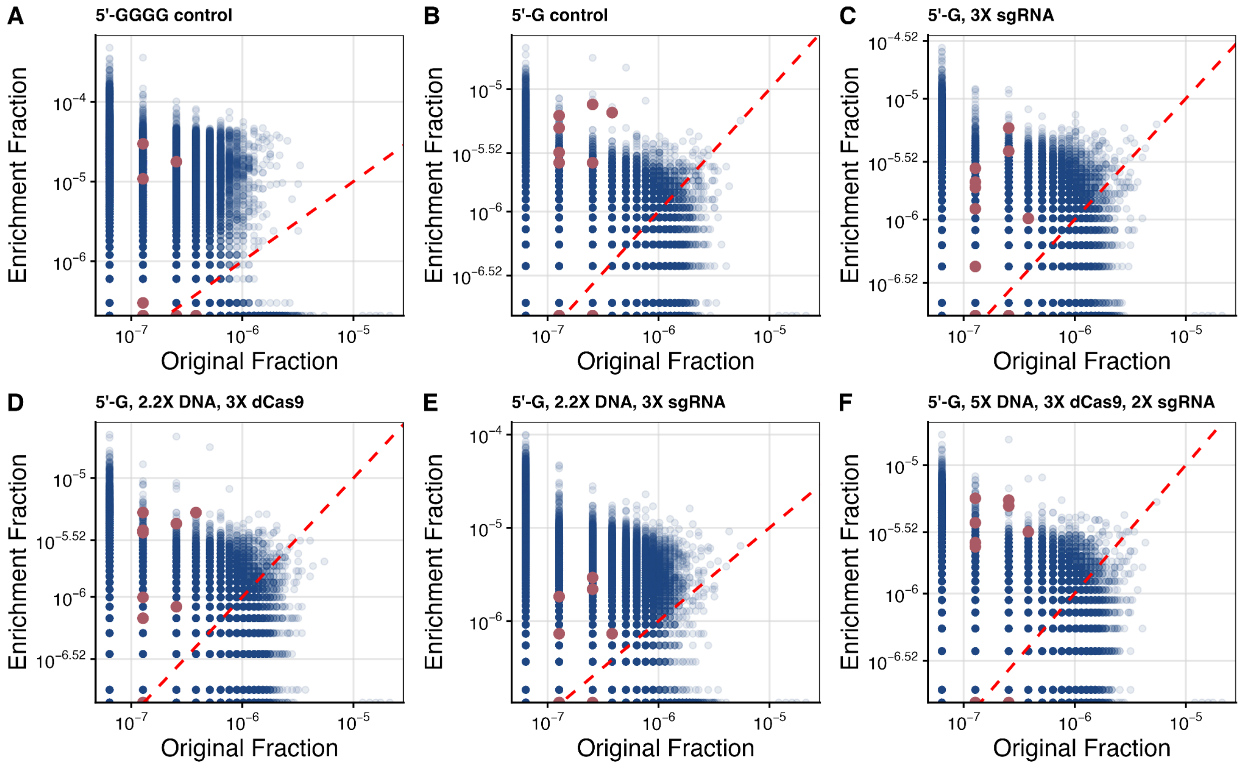

**Figure S15. Barcode distributions before and after 12-plex enrichment of a 384-gene DHFR library (library S4) with varying amounts of input DNA, sgRNAs, and dCas9.** Scatter plots compare the fractional abundance of barcodes in the 384-gene DHFR library (S4) before (original) and after enrichment. Blue dots represent non-target barcodes, while magenta dots represent 12 targeted barcodes enriched under the experimental conditions listed above each plot. All enrichments used 5′ G sgRNAs, except for panel **A**, which used 5′ GGGG sgRNAs. The red dashed unity line indicates equal representation before and after enrichment, serving as a reference for assessing enrichment. The following conditions were evaluated: **A.** BAR-CAT v1.0 control enrichment using 5′ GGGG sgRNAs, **B.** BAR-CAT v1.0 control enrichment using 5′ G sgRNAs, **C.** Increasing sgRNAs by 3x (5’ G) **D.** Increasing input DNA by 2.2x (5’ G) and dCas9 by 3x , **E.** Increasing input DNA by 2.2x and sgRNAs by 3x (5’ G) **F.** Increasing enrichment volume by 2x, sgRNAs by 2x, input DNA by 5x, and dCas9 by 3x.

**Table S1.** Compositions of wash buffers tested for streptavidin bead washing stringency. The qPCR results for bead wash stringency are in **Fig. S1B**.

| Wash buffer | Buffer ID | Buffer Composition |
|---|---|---|
| Immobilization buffer (IB) | 1 | **1 mM DTT,** 10 mM Tris-HCl, 1 mM EDTA●$Na_{2,}$ pH 8.0 |
| 2X binding and wash buffer (2X B&W) | 2 | **2M NaCl**, 1 mM EDTA, 10mM Tris-HCl pH 7.4 |
| 2X B&W, 10% NP-40 | 3 | **2M NaCl**, **10% NP-40,** 1mM EDTA, 10mM Tris-HCl pH 7.4 |
| 1X TE, 10% NP-40 | 4 | **10% NP-40,**1 mM EDTA, 10mM Tris-HCl pH 7.4 |

**Table S2.** Summary of all PCR primers used for Illumina library preparation, BAR-CAT enrichment, amplicon sequencing, and other PCR applications.

| DNA Primer Name | Primer Type | Sequence | Modifications | Description |
|---|---|---|---|---|
| RFP_KpnI_FWD_Biotin_NV | Forward | CGTGGTACCTTATTTGTACAGCTCATC | 5’ biotin | Biotinylation and amplification of *mcherry* gene insert. |
| RFP_Ndel_REV_Biotin_NV | Reverse | CGTGGTACCTTATTTGTACAGCTCATC | 5’ biotin | Biotinylation and amplification of *mcherry* gene insert. |
| skpp504F | Forward | ATCGGGGATGGTAACTAACG | 5’ biotin | Biotinylation and amplification of library S4 DHFR gene inserts. |
| skpp504R | Reverse | ATAGCTGATTGTCCGTTGGT | 5’ biotin | Biotinylation and amplification of library S4 DHFR gene inserts. |
| P5_FWD_Biotin_NV | Forward | AATGATACGGCGACCACCGAGATCTACAC | 5’ biotin | Biotinylation and amplification of library S2 DHFR gene inserts. |
| P7_REV_Biotin_NV | Reverse | ATCTCGTATGCCGTCTTCTGCTTG | 5’ biotin | Biotinylation and amplification of library S2 DHFR gene inserts. |

| | | | | |
|---|---|---|---|---|
| mi7_FWD_Amp_NV | Forward | AATGATACGGCGACCACCGAGATCTACACCTGGCCGTAAGGTACCTAAGTGTCGCTGCCGAACAGG | N/A | Used for addition of the P5 Illumina adapter for sequencing pEVBC3-RFP barcodes; also used for post-enrichment amplification in BAR-CAT v0.1, v0.2. |
| mi7_REV_Amp4_NV | Reverse | CAAGCAGAAGACGGCATACGAGATTCGCCTTACGTTGGCCACCCAAGTCATTCTGAGAATAGTGTATGCGGCGACCG | N/A | Addition of P7 Illumina adapter and Illumina. N701 index 1 for sequencing pEVBC3-RFP barcodes. |
| Mi7_R1_NV | Illumina read 1 | CTGGCCGTAAGGTACCTAAGTGTCGCTGCCGAACAGG | N/A | Illumina sequencing pEVBC3-RFP |
| Mi7_R2_NV | Illumina read 2 | CGAAAAGGAATTCTGCGACGTGACGTCGTGCGCTCTTCTAGC | N/A | Illumina sequencing pEVBC3-RFP |
| Mi7_Rindex | Illumina index read | GCGCAAACTATTAACTGGCGAACTACTTACTCTAGCTTCCCGGCCCAGTAAC | N/A | Illumina sequencing pEVBC3-RFP |
| Mi8_REV_Amp4_Index2_NV | Reverse | CAAGCAGAAGACGGCATACGAGATCTAGTACGCGTTGGCCACCCAAGTCATTCTGAGAATAGTGTATGCGGCGACCG | N/A | Illumina primer used for post-enrichment amplification in BAR-CAT v0.1, 0.2. |
| mi9_FWD_Amp_NV | Forward | AATGATACGGCGACCACCGAGATCTACACGCCGCCGAACGAC | N/A | Used for addition of the P5 Illumina adapter for sequencing |

| | | | | |
|---|---|---|---|---|
| | | CGAGCGCAGCCATA | | pEVBC8-DHFR (libraries S2 and S4); also used for post-enrichment amplification in BAR-CAT v0.3, v1.0 |
| Mi9_REV_amp_NV | Reverse | CAAGCAGAAGACGGCATACGAGATTCGCCTTACGAAAAGGAATTCTGCGACGTGACGTCGTG | N/A | Illumina primer used for post-enrichment amplification in BAR-CAT v0.3, v1.0. |
| mi9_REV_Amp_Index2_NV | Reverse | CAAGCAGAAGACGGCATACGAGATCTAGTACGCGAAAAGGAATTCTGCGACGTGACGTCGTG | N/A | Addition of P7 Illumina adapter and Illumina N702 index 2 for sequencing pEVBC8-DHFR lib S2 |
| mi9_REV_Amp_Index5_NV | Reverse | CAAGCAGAAGACGGCATACGAGATAGGAGTCCCGAAAAGGAATTCTGCGACGTGACGTCGTG | N/A | Addition of P7 Illumina adapter and Illumina N705 index 5 for sequencing pEVBC8-DHFR lib S4 |
| Mi9_R1_NV | Illumina read 1 | GCCGCCGAACGACCGAGCGCAGCCATATG | N/A | Illumina sequencing pEVBC8-DHFR libs S2 and S4 |
| Mi9_R2_NV | Illumina read 2 | CGAAAAGGAATTCTGCGACGTGACGTCGTGCGCTCTTCTCCT | N/A | Illumina sequencing pEVBC8-DHFR libs S2 and S4 |
| Mi9_EVBC8_Rindex | Illumina index read | AGGAGAAGAGCGCACGACGTCACGTCGCAGAATTCCTTTTCG | N/A | Illumina sequencing pEVBC8-DHFR libs S2 and S4 |

| | | | | |
|---|---|---|---|---|
| qPCR_RFP_NV_FWD | Forward | GTATGTTAAACACCCAGCGG | N/A | qPCR amplification of *mcherry* from pEVBC3-RFP to evaluate bead wash stringency |
| qPCR_RFP_NV_REV | Reverse | GTCACGACACCACCATCTTC | N/A | qPCR amplification of *mcherry* from pEVBC3-RFP to evaluate bead wash stringency |

**Table S3**. Summary of PCR reactions used to build the pEVBC3 and pEVBC8 barcoding plasmids via round-the-horn PCR, including primers (from **Table S4**), templates, and resulting products. All PCRs were performed at an annealing temperature of 72 °C using Q5® High-Fidelity DNA Polymerase (New England Biolabs).

| Barcoding Plasmid | PCR Step | PCR Template | Forward Primer Name | Reverse Primer Name | PCR Product | PCR Cycle Number |
|---|---|---|---|---|---|---|
| pEVBC3 | PCR 1 | Linearized pEVBC1 | EVBC3_FWD1 | EVBC3_REV1 | pEVBC3 backbone | 35 |
| pEVBC3 | PCR 2 | pEVBC3 backbone (PCR 1 product) | EVBC3_FWD2 | EVBC3_REV2 | Barcoded pEVBC3 backbone | 5 |
| pEVBC3 | PCR 3 | barcoded pEVBC3 backbone (PCR 2 product) | EVBC3_FWD3RE | EVBC3_REV2 | Amplified and barcoded pEVBC3 backbone | 15 |
| pEVBC8 | PCR 1 | Linearized pEVBC3 | EVBC8_FWD1 | EVBC3_REV1 | pEVBC8 backbone | 35 |
| pEVBC8 | PCR 2 | pEVBC8 backbone | EVBC8_FWD2 | EVBC3_REV2 | Barcoded pEVBC8 backbone | 5 |
| pEVBC8 | PCR 3 | Barcoded pEVBC8 | EVBC3_FWD3RE | EVBC3_REV2 | Amplified and barcoded | 15 |

| | | | | | |
|---|---|---|---|---|---|
| | | backbone | | | pEVBC8 backbone |

**Table S4.** PCR primer sequences used to build barcoding plasmids, pEVBC3 and pEVBC8, by using three sequential PCR steps as part of a PCR-round-the-horn strategy.

| DNA Primer Name | Forward or Reverse | Sequence | Modifications |
|---|---|---|---|
| EVBC3_FWD1 | Forward | GCTAGAAGAGCGCACGACGTCACGTCGCAGAATTCCTTTTCGGGGAAATGTGCGCGGAAC | N/A |
| EVBC3_REV1 | Reverse | GCCGTCATATGGCTGCGCTCGGTCGTTCGGCTGCGGCGAGGAATTCCGGTATCAGCTCACTCAAAGGCGGTAATACG | N/A |
| EVBC3_FWD2 | Forward | GTGGGTACCTAAGTGTCGCTGCCGAACAGGNNNNNNNNNNNNNNNNNNNNGCTAGAAGAGCGCACGACGTCACGTC | N/A |
| EVBC3_REV2 | Reverse | GCCGTCATATGGCTGCGCTCGG | 5’ biotin |
| EVBC3_FWD3RE | Forward | GTGGGTACCTAAGTGTCGCTGCCGAACAG | 5’ biotin |
| EVBC8_FWD1 | Forward | AGGAGAAGAGCGCACGACGTCACGTCGCAGAATTCCTTTTCGGGGAAATGTGCGCGGAAC | N/A |
| EVBC8_FWD2 | Forward | GTGGGTACCTAAGTGTCGCTGCCGAACAGCNNNNNNNNNNNNNNNNNNNNAGGAGAAGAGCGCACGACGTCACGTC | N/A |

**dCas9 Enrichment Master Protocol**

1. **Prepare ribonucleoprotein complexes (RNPs)**. RNPs consist of the dCas9 protein complexed to single guide RNAs (sgRNA). sgRNAs provide targeted specificity once dCas9 recognizes a three-nucleotide proximal adjacent motif (PAM) on the DNA non-target strand. Prepare all dilutions and RNPs in PCR tubes (0.2 mL) on a metal PCR cooling rack on ice.
   a. Make the following dilutions before preparing RNPs
      i. Dilute sgRNA pool to 300 nM (~11.3 ng/µL for sgRNAs that are 100 bases in length) with nuclease-free water.
      ii. Dilute dCas9-3xFLAG-Biotin Protein (Sigma-Aldrich cat: DCAS9PROT-50UG) to 1 uM with dilution buffer following manufacturer instructions.
   b. Prepare RNPs by adding reagents according to the following table below in the listed order.

| Condition | Enrichment RNPs | No template control (NTC) RNPs |
|---|---|---|
| RNP formation ratio (dCas9:sgRNAs) | 3:1 | 3:1 |
| Component | | |
| Murine RNase inhibitor (NEB cat: M0314S) | 0.75 µL | 0.75 µL |
| Nuclease-free water to fill | 19.25 µL | 19.25 µL |
| 10XNEBuffer r3.1 (NEB cat: B6003S) | 3 µL | 3 µL |
| Pooled sgRNAs (300 nM) | 3 µL | 3 µL |
| Biotinylated dCas9 (1 µM) | 1 µL | 1 µL |
| Total vol: | 27 µL | 27 µL |

   c. Mix thoroughly and pulse-spin in a microfuge.
   d. Pre-incubate samples at 25°C within a thermocycler for 10 minutes
   e. Incubate samples at 37°C within a thermocycler for 10 minutes

2. **Prepare enrichment reactions by adding library DNA to RNPs.**
   a. Make sure that library DNA is sufficiently concentrated to add to RNPs. Add a total of 500 ng of DNA to each enrichment, making sure to add no more than 3 µl.
      a. In this case, I used EVBC3-RFP (~238 ng/µL) as the template for enrichment. This library contains the mCherry gene flanked by thousands of unique 20-mer barcodes.
   b. Prepare reactions according to the table below

| Condition | Enrichment | No template control (NTC) |
|---|---|---|

| | | |
|---|---|---|
| RNPs (**from step 1b**) | 27 µL | 27 µL |
| EVBC3-RFP scape DNA (238 ng/ul) | 2.1 µL | N/A |
| Nuclease-free water to fill | 0.9 µL | 3 µL |
| **Final vol:** | **30** µL | **30 µL** |

c. Mix thoroughly and pulse-spin reactions in a microfuge.
d. Incubate at 37°C for 15 minutes in a thermocycler (hold at 4°C if you can't remove samples immediately)

3. **Perform pulldown of biotinylated dCas9 with streptavidin-coated magnetic beads (NEB Cat: S1420S).** This step enables pull-up of dCas9 bound to targeted DNA barcodes.
   a. Clean beads according to the manufacturer's instructions.
      i. Dilute 2X B&W buffer (10mM Tris-HCl, 1mM EDTA•$Na_2$, pH 7.5, 2M NaCl) 2-fold with nanopure water to make 1X B&W.
      ii. Clean 5 µl of beads per enrichment reaction (including NTC).
         1. Vortex the stock bottle of beads for 30 seconds and then withdraw the total volume of beads you are cleaning into a 1.5 mL tube.
         2. Add 1 mL of 1x B&W buffer to the volume of beads that you intend to clean
         3. Place tube on magnetic rack for 1 minute and discard supernatant
         4. Resuspend beads in the same volume of 1x B&W buffer that you originally started with
         5. Repeat steps 3 and 4 three times.
         6. After final wash, resuspend clean beads in 2x B&W buffer, adding 2x the volume that you started with
   b. Add 10 µL of cleaned beads to nuclease-free 1.5 mL tubes.
   c. Add each 30 ul enrichment reaction (including NTC) to beads in 1.5 mL tubes immediately following enrichment incubation. Total volume should be ~40 µl.
   d. Shake beads (1700 rpm) in thermomixer at 37°C for 30 min.

4. **Wash streptavidin-coated magnetic beads bound to biotinylated dCas9 to remove non-target DNA prior to PCR amplification and sequencing.**
   a. Spin down enrichments containing beads (from step 3d) and transfer to fresh 5 mL tubes
   b. Wash beads in enrichment and NTC samples nine times with 2 mL 2X B&W buffer.
      i. Place 5 mL tubes on magnetic rack that can contain 15 mL tubes for 1 minute
      ii. Discard supernatant and proceed with the next wash
      iii. Transfer samples to fresh 5 mL tubes right before the sixth wash.
   c. After washing, resuspend enrichment samples in 50 µL nuclease-free water. Resuspend NTC in 20 µL nuclease-free water.

        i. Vortex beads (2700 rpm) and then spin them down using a centrifuge with a rotor for 5 mL tubes.
        ii. Samples can be stored at 4°C overnight

5. **Treat samples with proteinase K (NEB Cat: P8107S) to denature dCas9 and release enriched DNA into the supernatant.**
    a. Transfer 50 µL of resuspended beads (from step 4c) to fresh 1.5 mL tubes prior to proteinase K treatment
    b. Add 1 µL proteinase K to enrichment samples
    c. Mix samples thoroughly and pulse-spin in a microfuge
    d. Incubate samples for 10 min at room temperature (~25°C) in thermomixer with shaking (1700 rpm).
    e. Place samples on a magnetic rack for one minute to separate beads from supernatant.
    f. Collect the supernatant and place it in a fresh 1.5 mL tube,
    g. Clean the supernatant using Monarch DNA clean-up kit.
        i. Use 100 µL of DNA binding buffer per sample (two times 50 µL sample volume)
        ii. Elute enrichment products with 20 µL of hot elution buffer in one 5 µg DNA clean-up column per sample (NEB Cat: T1034-2).
    h. Use clean DNA for downstream PCR.
6. **Prepare quantitative polymerase chain reactions (qPCR) of enrichment samples to determine the number of cycles needed to amplify enrichment DNA products and avoid overamplification**.
    a. Prepare qPCRs according to the table below with Q5 2x master mix (NEB cat: M0492) and 100X thiazole green (Biotium cat: 40086). Prepare duplicates for all enrichment samples, including the NTC. Scale up master mix as needed.

| Component | 1x reaction (50 µL total volume) |
|---|---|
| Water to fill | 17.5 µL |
| Q5 2X master mix | 25 µL |
| Mi9_FWD_Amp_NV (10 uM) | 2.50 µL |
| Mi9_REV_amp_NV (index 1) (10 uM) | 2.50 µL |
| Enrichments/NTC (from step 53) | 2 µL (add after aliquoting master mix) |
| Thiazole green (100X) | 0.5 µL |

    b. Aliquot 48 µL of master mix/rxn and add templates last.
    c. Run according to PCR parameters (table below)

| Cycles | Step | Temp | Time |
|---|---|---|---|
| 1 | Denaturation | 98°C | 30 s |
| 60 | Denaturation | 98°C | 10 s |
| | Annealing | 72°C | 30 s |

| | | | |
|---|---|---|---|
| | Extension | 72°C | 30 s |

d. Choose the lowest number of cycles to bulk amplify enriched DNA.

7. **Bulk-amplify the enrichment products for sequencing by performing PCR**.
    a. Prepare a 7X master mixes per enrichment sample and 2X PCRs for the NTC using the table below.

| Component | 1x reaction (50 µL reactions) |
|---|---|
| Water to fill | 18 µL |
| Q5 2X master mix | 25 µL |
| Mi9_FWD_Amp_NV (10 uM) | 2.50 µL |
| Mi9_REV_amp_NV (index 1) (10 uM) | 2.50 µL |
| Enrichments/NTC (from step 53) | 2 µL |

    b. Aliquot 50 µl of each master mix into PCR tubes. Run according to PCR parameters in the table below

| Cycles | Step | Temp | Time |
|---|---|---|---|
| 1 | Denaturation | 98°C | 30 s |
| Use cycle numbers determined in step 6 | Denaturation | 98°C | 10 s |
| | Annealing | 72°C | 30 s |
| | Extension | 72°C | 30 s |
| 1 | Final extension | 72°C | 2 min |
| 1 | hold | 12°C | infinite |

    c. Use Monarch DNA clean-up kit to clean PCR products.
        i. Pool samples and clean using DNA binding buffer equivalent to 2X the volume of pooled samples.
        ii. Elute each enrichment sample using a Monarch miniprep DNA clean-up column (NEB Cat: T1017-2) and 30 µL of hot elution buffer. Elute the NTC with a 5 µg DNA clean-up column and 8 µl of hot elution buffer

8. **Size-select enriched DNA PCR products to eliminate off-target PCR products.**
    a. Prepare a 2% Agarose gel (TAE) to run and gel extract enrichment products
    b. Run gel at 115 V for 1 hour
    c. Stain with either SYBR safe (APExBIO Cat: A8743) or SYBR gold (ThermoFisher Cat: S11494) depending on the concentration of DNA you are loading into the gel.
    d. Size-select correct PCR products by cutting correctly sized DNA bands with a razor.
    e. Clean size-selected DNA using the Monarch gel extraction DNA clean-up kit.

a. Use one miniprep column to clean each sample and elute with 30 µL hot elution buffer.

9. Submit 200-300 ng of DNA from step 8 for nanopore sequencing to evaluate enriched barcodes.

**Supplementary methods**

***Examining the Impact of Magnetic Bead Washes on Removal of Non-enriched DNA***

Before enriching the pilot library (pEVBC3-RFP) with BAR-CAT 0.1, the effect of magnetic bead washes on DNA recovery was assessed to establish optimal washing conditions. Due to the unconfirmed functionality of biotinylated dCas9 for enrichment, EnGen® Spy dCas9 (SNAP-tag®) (NEB) and SNAP-Capture Magnetic Beads (NEB) were used for CRISPR enrichments. Each enrichment reaction, except the no library DNA control, included 50 ng of the pilot library (pEVBC3-RFP). Following enrichment, beads were washed 15 times with 1 mL of immobilization buffer (IB) (**Table S1**) according to the NEB protocol for SNAP-capture magnetic beads. Supernatant from washes 3, 6, 9, 12, and 15, including the no template control, was collected.

***Quantitative PCR (qPCR) and Agarose Gel Electrophoresis Analysis***

Quantitative PCR (qPCR) was performed to compare Cq values across washes. Reactions were prepared using the supernatants collected from dCas9 capture bead washes, with beads from wash 15 included as a positive control. Each qPCR reaction contained 3.75 µl of wash supernatant or SNAP-tag dCas9 capture beads, 1.25 µl each of 10 µM forward and reverse primers, 0.25 µl of 100X Biotium Thiazole Green (Thermo Fisher Scientific), 12.5 µl of 2X Q5 High-Fidelity Master Mix (NEB), and nuclease-free water to a total volume of 25 µl, following the Illumina-indexed primer amplification protocol for MiSeq of the pEVBC3-RFP library (see ***Indexed amplification of the barcoded rfp gene library for MiSeq*** in **Methods**). Since qPCR Cq values were uninformative, the 586 bp amplified products were purified using Monarch® 5 µg DNA Cleanup Columns (NEB). Purified DNA was visualized with SYBR™ Safe DNA Gel Stain (Thermo Fisher) following agarose gel electrophoresis on a 2% gel, using a 100 bp ladder (NEB) as a reference. DNA band intensities were quantified using ImageJ (1).

***qPCR Evaluation of Buffer Composition on the Stringency of Streptavidin-Coated Magnetic Bead Washes***

Prior to the development of BAR-CAT 0.1, the ability of biotinylated dCas9 to mediate enrichment was unverified. To assess and improve wash stringency, four buffer compositions compatible with streptavidin-coated magnetic beads were evaluated (see **Table S1**). Buffers containing high salt (2 M NaCl) or an anionic detergent (10% NP-40) were predicted to enhance wash stringency.

Streptavidin-coated magnetic beads (Sigma) were equilibrated according to the manufacturer's instructions, then resuspended in 2X B&W buffer (2 M NaCl, 1 mM EDTA, 10 mM Tris-HCl, pH 7.4) at twice the original volume. Ten microliters of equilibrated beads were transferred to individual 1.5 mL tubes.

Each bead aliquot received 1 µL of 10 ng/µL pEVBC3-RFP plasmid containing barcode 12 (BC12). Beads were incubated for 30 minutes at 37 °C with shaking at 1700 rpm. Following incubation, beads were washed either six times (for RT and 30 °C conditions) or three times (for RT) with 50 µL of the corresponding buffer. Washes were performed using magnetic separation or vortexing followed by magnetic separation between steps. After the final wash, beads were resuspended in 10 µL of nuclease-free water.

To assess wash efficiency, qPCR was performed to quantify the remaining pEVBC3-RFP BC12 template in the final wash supernatants. qPCRs for each wash condition were prepared using either 3 µL or 3.3 µL of final wash supernatant as template. Each reaction included 0.4 µL of 10 µM forward (qPCR_RFP_NV_FWD) and reverse (qPCR_RFP_NV_REV) primers (**Table S2**), 10 µL of KAPA SYBR Fast 2X Master Mix (Roche), and nuclease-free water to a final volume of 20 µL. A positive control (10 ng pEVBC3-RFP BC12) and no-template controls were included.
